# Two paralogous PHD finger proteins participate in *Paramecium tetraurelia*’s natural genome editing

**DOI:** 10.1101/2024.01.23.576875

**Authors:** Lilia Häußermann, Aditi Singh, Estienne C. Swart

**Affiliations:** Max Planck Institute for Biology, Max-Planck-Ring 5, 72076, Tübingen, Germany

**Keywords:** genome reorganization, PHD finger proteins, small RNAs

## Abstract

The unicellular eukaryote *Paramecium tetraurelia* contains functionally distinct nuclei: germline micronuclei (MICs) and a somatic macronucleus (MAC). During sexual reproduction, the MIC genome is reorganized into a new MAC genome and the old MAC is lost. Almost 45,000 unique Internal Eliminated Sequences (IESs) distributed throughout the genome require precise excision to guarantee a functional new MAC genome. Here, we characterize a pair of paralogous PHD finger proteins involved in DNA elimination. DevPF1, the early-expressed paralog, is present in only some of the gametic and post-zygotic nuclei during meiosis. Both DevPF1 and DevPF2 localize in the new developing MACs, where IESs excision occurs. In *DevPF2* knockdown (KD) long IESs are preferentially retained and late-expressed small RNAs decrease; no length preference for retained IESs was observed in *DevPF1*-KD and development-specific small RNAs were abolished. The expression of at least two genes from the new MAC with roles in genome reorganization seems to be influenced by *DevPF1-* and *DevPF2*-KD. Thus, both PHD fingers are crucial for new MAC genome development, with distinct functions, potentially via regulation of non-coding and coding transcription in the MICs and new MACs.

## Introduction

A unique feature shared by all ciliates is the presence of nuclear dimorphism. In *Paramecium tetraurelia* (henceforth *Paramecium*) the two micronuclei (MICs) resemble the germline of multicellular organisms, being transcriptionally silent throughout most of the life cycle and generating haploid nuclei during meiosis that develop and give rise to all nuclei in the subsequent generation. Also similar to the multicellular soma, the macronucleus (MAC) is optimized for most gene expression, and originates from a MIC copy. The old MAC is fragmented during sexual division and subsequently diluted across cell divisions, with the new MAC completely taking over somatic expression. The development from the MIC genome to the MAC genome in *Paramecium* is a natural form of genome editing that requires extensive reorganization, including genome amplification (∼800n), chromosome fragmentation and the elimination of about 25% of the sequence content (Arnaiz et al., 2012; Guérin et al., 2017). These MIC genome-specific sequences comprise repeats, transposable elements and Internal Eliminated Sequences (IESs).

In contrast to other elimination events, IES elimination requires precise excision in *Paramecium*. Precise IES excision is not characteristic of all ciliates. Notably, in *Paramecium*’s oligohymenophorean relative *Tetrahymena*, IESs are predominantly imprecisely excised and only tolerated in intergenic regions (Hamilton et al., 2016). The roughly 45,000 IESs in *Paramecium* are scattered throughout the genome in both non-coding and coding regions and vary from tens to thousands of base pairs in length (Arnaiz et al., 2012). Since the coding density of the *Paramecium* MAC genome is high, most IESs are intragenic (Arnaiz et al., 2012). *Paramecium* IESs are flanked by conserved 5’-TA-3’ dinucleotides (Klobutcher & Herrick, 1995) and excised by PiggyMAC (Pgm). Pgm is a domesticated transposase derived from PiggyBac transposases (Baudry et al., 2009) like the excisase responsible for IES excision in *Tetrahymena* (Cheng et al., 2010). The weakly conserved ∼5 bp long inverted repeats at *Paramecium* IES ends (Klobutcher & Herrick, 1995) fail to provide enough specificity for reliable Pgm recruitment (Arnaiz et al., 2012). This suggests that other factors are needed for precise IES targeting.

The targeting of MIC-specific sequences for elimination is thought to be assisted by small non-coding RNAs, first characterized in *Tetrahymena* (Chalker & Yao, 2001; Mochizuki et al., 2002). Like *Tetrahymena*, the biogenesis of the 25 nucleotide (nt) scan RNAs (scnRNAs) occurs during meiosis in the *Paramecium* MICs. Bidirectional non-coding transcription of the MIC genome is thought to be initiated by the putative transcription elongation factor Spt5m (Gruchota et al., 2017) and followed by the cleavage of long double-stranded RNA (dsRNA) by the closely related Dicer-like protein paralogs Dcl2 and Dcl3 (Hoehener et al., 2018; Lepère et al., 2009; Sandoval et al., 2014). Argonaute/Piwi proteins Ptiwi01/09 (also close paralogs) process the resulting short dsRNAs, removing one of the two strands, and stabilize single-stranded scnRNAs throughout the selection process in the parental MAC and targeting of MIC-specific sequences in the new MACs (Bouhouche et al., 2011; Furrer et al., 2017). In the parental MAC, Gtsf1 was recently proposed to promote ubiquitination and subsequent degradation of the Ptiwi01/09 complexes harboring MAC-matching scnRNAs (Charmant et al., 2023; Wang et al., 2023). In the new MACs, the putative transcription elongation factor TFIIS4 was proposed to promote non-coding transcription required for scanning the developing genome (Maliszewska-Olejniczak et al., 2015).

In *Tetrahymena*, H3K9 and H3K27 methylation precede IES excision (Y. Liu et al., 2007; Taverna et al., 2002) and it was shown in *Paramecium* that development-specific H3K9me3 and H3K27me3 histone mark deposition by the PRC2 complex depends on scnRNAs and is essential for the elimination of transposons and IESs (Frapporti et al., 2019; Ignarski et al., 2014; Lhuillier-Akakpo et al., 2014; Miró-Pina et al., 2022; Wang et al., 2022). We recently showed that the ISWI1 chromatin remodeling complex is necessary for IES excision precision and Ptiwi01/09 co-immunoprecipitated with ISWI1 in a crosslinked treatment (Singh et al., 2022, 2023). After the initial onset of IES excision, additional single-stranded sRNAs, iesRNAs, ranging in size from ∼26 to 30 bp, are produced by Dcl5 from excised IES fragments and stabilized on Ptiwi10/11 (Furrer et al., 2017; Sandoval et al., 2014). iesRNAs were proposed to participate in a positive feedback loop for the efficient removal of all IES copies (Sandoval et al., 2014). Nevertheless, only a fraction of IES excision appears to depend on scnRNAs or iesRNAs (Nowacki et al., 2005; Sandoval et al., 2014).

Despite the knowledge gained in the past decades, the picture of IES excision is far from complete. To identify novel genes involved in IES excision, we examined proteins potentially associated with ISWI1, a chromatin remodeler we recently showed to facilitate precise IES excision (Singh et al., 2022).

## Results

### Identification of a novel protein involved in IES excision

Recently, we reported evidence supporting the formation of a protein complex involving ISWI1 and the ICOP proteins (Singh et al., 2023). We conducted an RNAi screen of additional genes that were unique in the ISWI1 co-immunoprecipitation (IP)-mass spectrometry (MS) and exhibited upregulation in a developmental gene expression time course from ParameciumDB (Arnaiz et al., 2017) (Fig. S1A).

In the screening, we sought phenotypic evidence for failed genome reorganization in the form of growth defects (assessed by survival tests), and substantial IES retention (assessed by IES retention PCRs). *ND7*, a gene involved in trichocyst discharge (Lefort-Tran et al., 1981), was used as a negative control as its silencing does not impair genome reorganization (Nowacki et al., 2005). *Nowa1*-KD, which affects the excision of scnRNA-dependent IESs (Nowacki et al., 2005), was used as a positive control. Candidate 2 (PTET.51.1.G0620188) displayed both IES retention and lethality in the new progeny, whereas candidate 1 (PTET.51.1.G0990120) showed high lethality without IES retention (Fig. S1B,C). Therefore, candidate 2 was selected for further investigations.

### DevPF2 and DevPF1 are paralogous PHD finger proteins

The *Paramecium aurelia* species complex, to which *P. tetraurelia* belongs, underwent multiple whole-genome duplications, with many closely related paralogs generated from the most recent of these (Sellis et al., 2021). The chosen candidate has a closely related paralog (PTET.51.1.G0240213) with which it shares 86.6% identity at both the nucleotide and amino acid levels. The paralog is upregulated during sexual development as well, although earlier (Fig. 1A). HMMER3 searches of the Pfam database (Finn et al., 2003) predicted two domains in both proteins: a PHD and a PHD zinc-finger-like domain (Fig. 1B,D,E).

**Figure 1:**
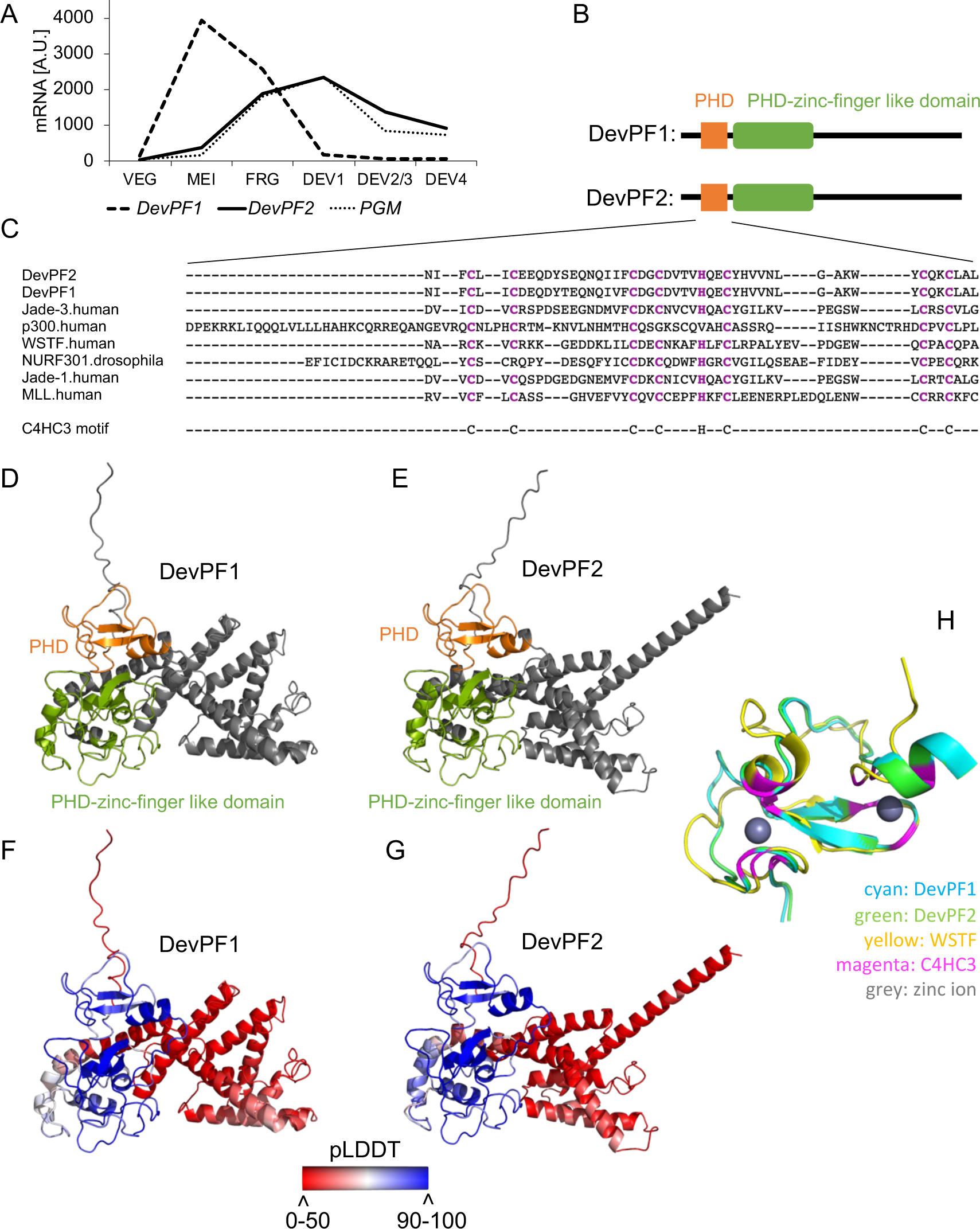
Features of the PHD finger proteins DevPF1 and DevPF2. (A) mRNA expression profiles for *DevPF1*, *DevPF2* and *PGM* during various developmental stages: VEG (vegetative growth), MEI (micronuclear meiosis and macronuclear fragmentation), FRG (∼50% of the population with fragmented maternal MACs), DEV1 (significant proportion with visible anlagen), DEV2/3 (majority with visible anlagen), DEV4 (majority with visible anlagen). Expression data retrieved from ParameciumDB (Arnaiz et al., 2017). (B) Schematic representation of predicted domain architecture for DevPF1 and DevPF2. (C) Multiple sequence alignment (Clustal Omega) of DevPF1 and DevPF2 amino acid sequence with PHD domains of published human and *Drosophila* PHD finger proteins. (D) to (F): Predicted protein structure (AlphaFold2) for DevPF1 and DevPF2, colored by domain (PHD: orange; PHD-zinc-finger-like domain: green) in (D) and (E), and by prediction confidence (pLDDT: predicted local distance difference test) in (F) and (G). (H) Structure predictions of DevPF1 and DevPF2 PHD domain superimposed with NMR structure of WSTF PHD domain (PDB accession number 1F62).

The highly conserved PHD domain has often been reported to mediate the interaction of nuclear proteins with histone modifications (Sanchez & Zhou, 2011), but other binding affinities have also been described (see Discussion). PHD domains possess a well-conserved motif consisting of eight cysteine and histidine residues (C4HC3) that coordinate two zinc ions, thereby providing it with structural stability. The presence of the C4HC3 motif in both paralogs was confirmed using a multiple sequence alignment with PHD domains from well-established PHD finger proteins from *Homo sapiens* and *Drosophila melanogaster* (Fig. 1C).

AlphaFold2 predicted the structures of both paralogs with high confidence for the domains (Fig. 1F,G). We compared the PHD predictions with the published structure of the WSTF (Williams Syndrome Transcription Factor) PHD finger (Pascual et al., 2000). WSTF, associated with the Williams Syndrome (Lu et al., 1998), is a subunit of the ISWI-containing chromatin remodeling complex WICH (Bozhenok et al., 2002). The superimposition confirmed the orientation of the eight C4HC3 residues in the DevPFs towards the two zinc ions (Fig. 1H), supporting the idea that both paralogs function as PHD finger proteins. Since they show development-specific upregulation (Fig. 1A), we named the paralogs development-specific PHD finger 1 (DevPF1; early-expressed paralog) and 2 (DevPF2; late-expressed paralog).

### DevPF1 and DevPF2 show distinct nuclear localization

To determine the localization of both paralogs, we injected DNA constructs encoding DevPF1 and 2 C-terminally tagged with green fluorescent protein (GFP) into MACs of vegetative paramecia. The cells were collected during *Paramecium* sexual development for confocal microscopy. The injected cultures displayed no growth defects compared to non-transformed cells (Fig. S2A). However, we observed variable numbers for gametic MICs (Figs 2, 3) and new MACs (Fig. S2C) in some cells, which has been observed frequently for transgenes (e.g, Nowa1-GFP fusion; (Nowacki et al., 2005)).

**Figure 2:**
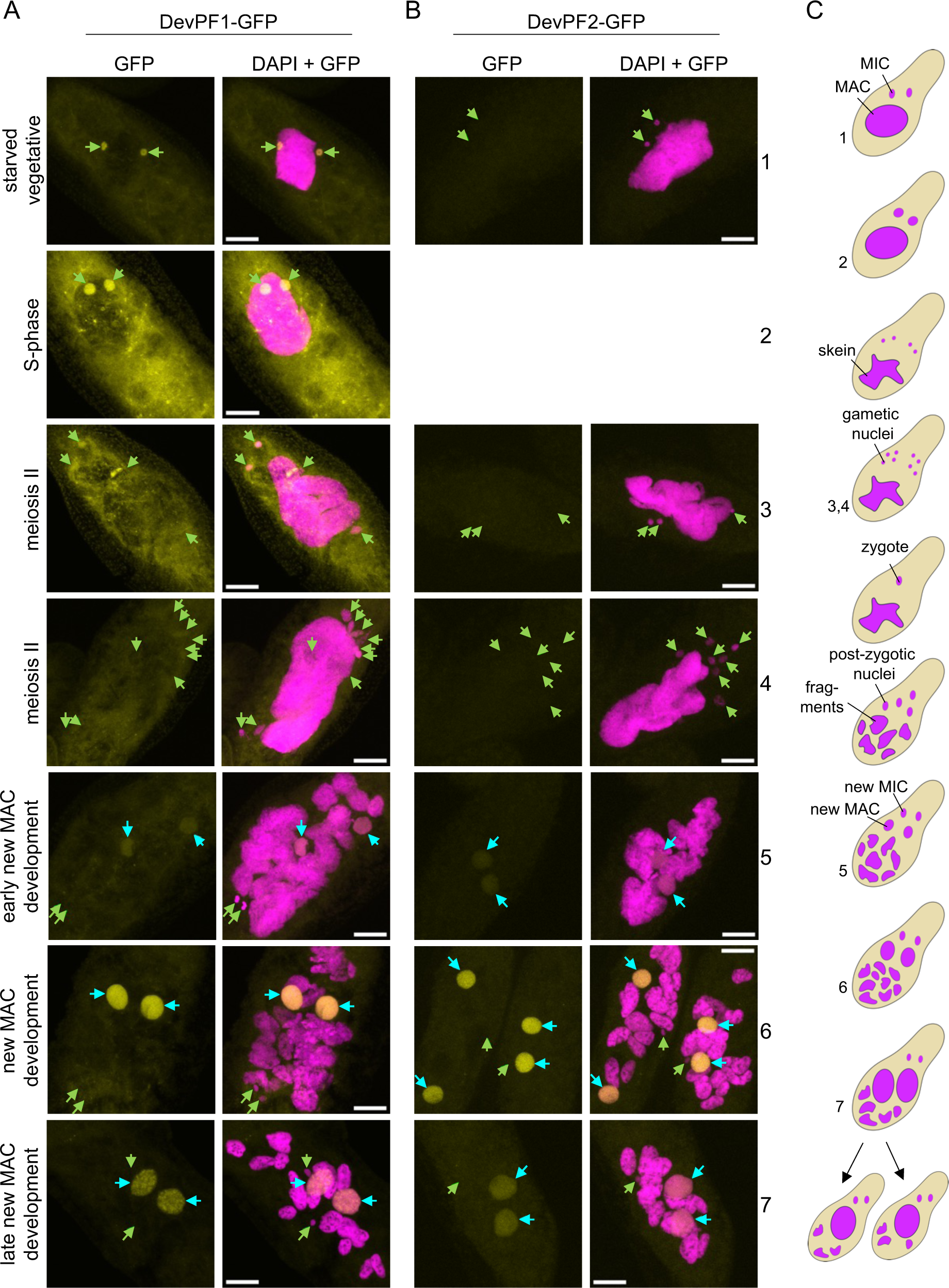
Subcellular localization of DevPF-GFP proteins. DevPF1-GFP (A) and DevPF2-GFP (B) localization at various developmental stages. DNA (stained with DAPI) in magenta. GFP signal in yellow. No image of DevPF2-GFP during S-phase was acquired. Green arrow: MIC. Cyan arrow: new MAC. Maximum intensity projections of multiple z-planes. Scale bar = 10 µm. (C) Schematic overview of nuclear morphology during sexual development, with corresponding cell stages in the images indicated by numbers.

Consistent with DevPF1’s early peak in mRNA expression from the developmental time course in ParameciumDB, DevPF1-GFP was expressed during the onset of sexual development, but not in vegetative cells with food vacuoles containing bacteria (Figs 2A, S2B). DevPF1-GFP was distributed throughout the cytoplasm and localized in both MICs before and during the S-phase of meiosis, when these nuclei swell (Fig. 2A). Throughout the subsequent meiotic divisions, DevPF1-GFP localized to only a few of the gametic MICs (Fig. 3A). Its micronuclear localization appeared independent of nuclear division as detected by the presence of the spindle apparatus (Fig. 3A,B). During post-zygotic mitotic divisions, DevPF1-GFP was observable in certain post-zygotic nuclei, but not in all of them (Fig. 3B). Later during development, DevPF1-GFP was present in the early new MACs and remained in the new MACs throughout development up to very late stages (Fig. 2A) despite the drop in its mRNA levels (Fig. 1B). During new MAC development there was also comparatively little cytoplasmic DevPF1-GFP compared to that during meiosis.

**Figure 3:**
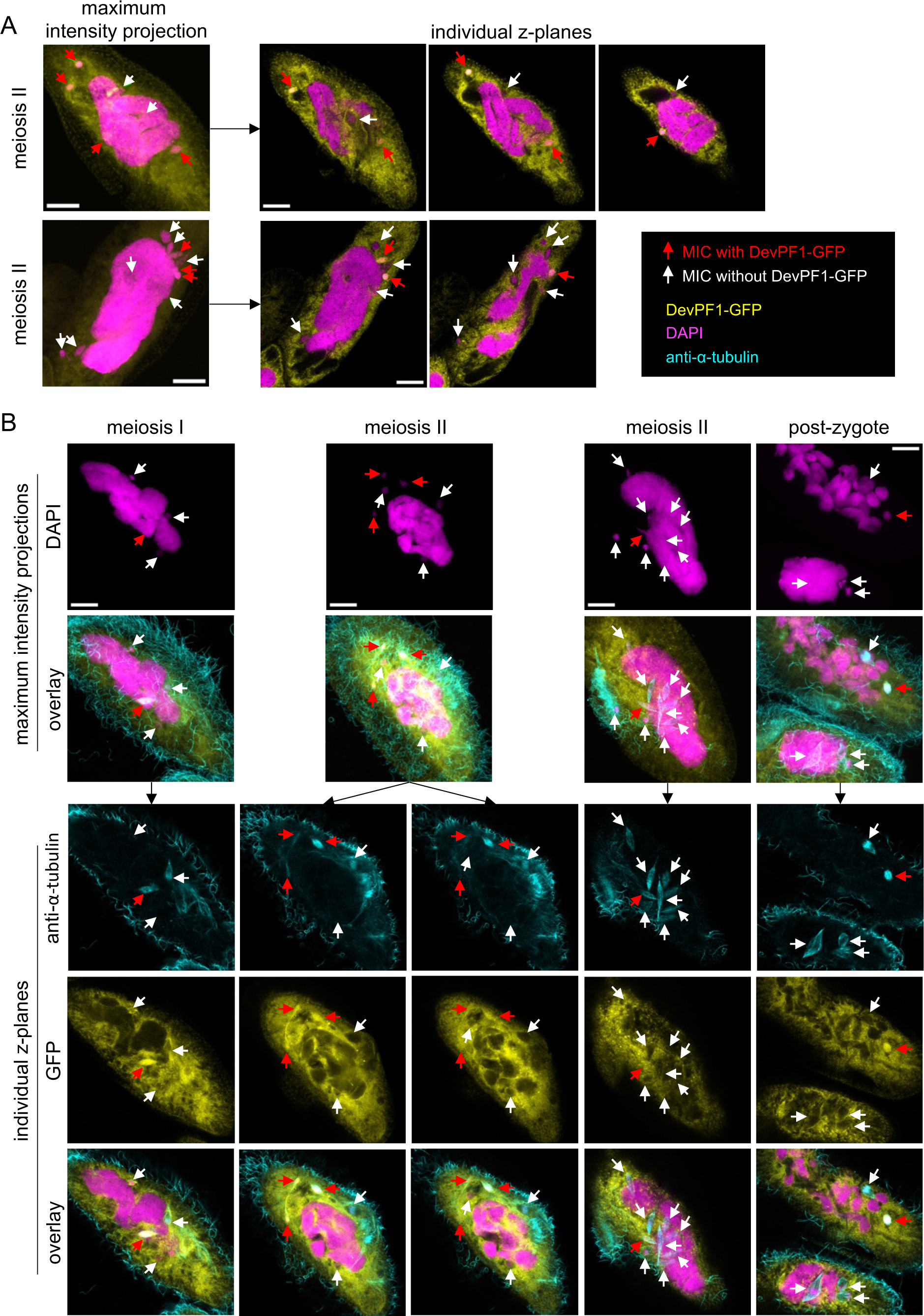
Selective DevPF1-GFP localization in *Paramecium* MICs. (A) Overlay of DAPI (DNA stain; pink) and GFP (yellow) signal in two DevPF1-GFP injected *Paramecium* cells during meiotic stages. Maximum intensity projections (left) and individual z-planes of the same stack (right). (B) DevPF1-GFP localization with visualization of nuclear spindle. DAPI (pink), GFP (yellow) and anti-α-tubulin staining (cyan). Maximum intensity projections (top) for DAPI and overlay (DAPI, GFP and anti-α-tubulin). Individual z-planes of the same stacks (bottom) for anti-α-tubulin, GFP and overlay. (A) and (B): Red arrows: MICs with DevPF1-GFP localization; White arrows: MICs without DevPF1-GFP localization. Scale bar = 10 µm.

Consistent with its mRNA expression profile, DevPF2-GFP emerged after the onset of new MAC development and localized within the new MACs, where it remained up to the late stages (Fig. 2B).

### Silencing constructs partially co-silence both paralogs

We utilized RNAi by feeding to investigate the influence of the DevPFs on IES excision. Two silencing regions were selected (a and b) on each *DevPF1* and *DevPF2* (Fig. 4A). Due to the lack of regions with sufficient specificity for either of the paralogs, co-silencing was predicted (see Methods). Hence, we first experimentally verified the possibility of co-silencing with mRNA and protein levels using silencing region a, since it exhibited less off-target hits.

**Figure 4:**
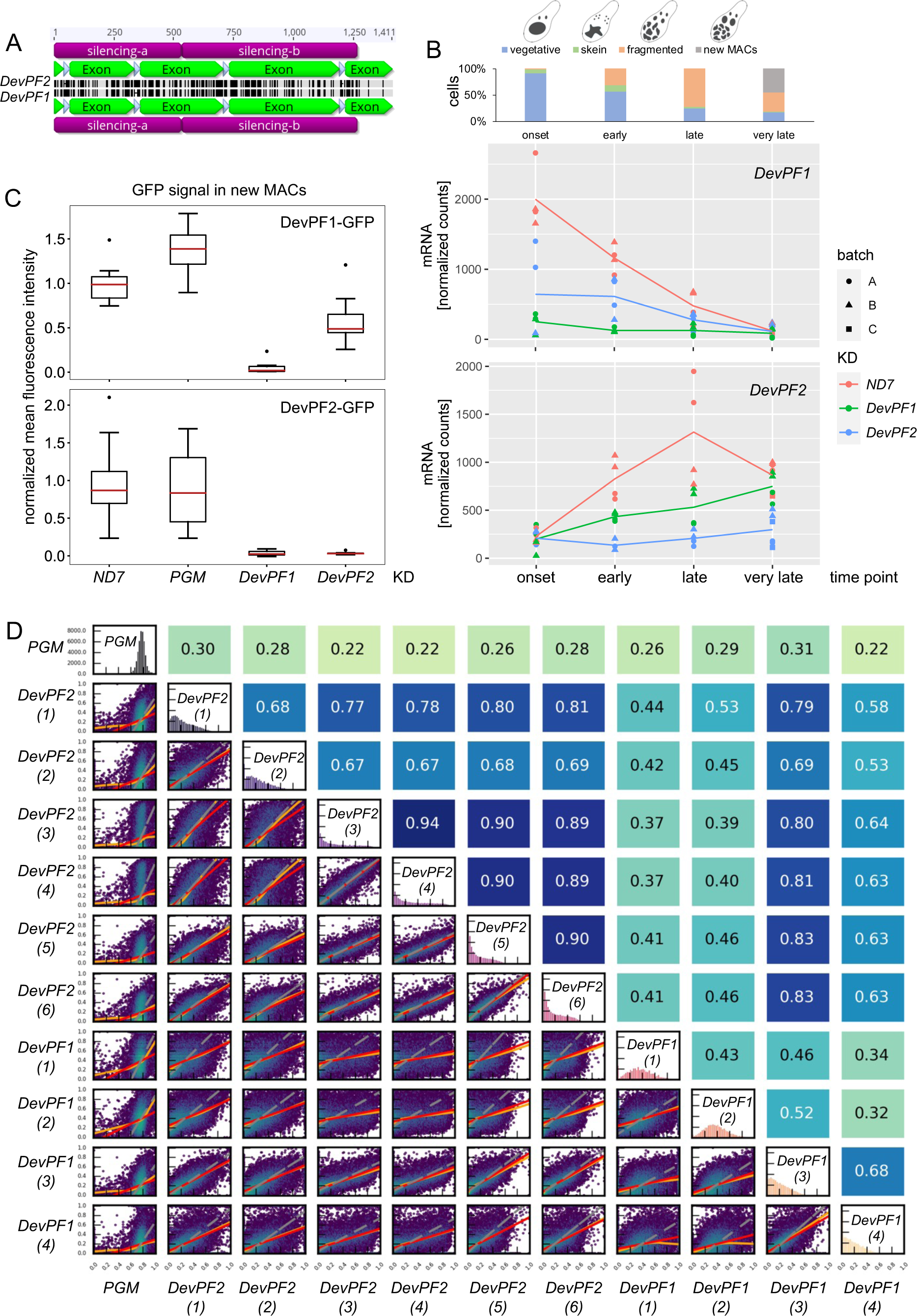
Co-silencing effects observed in *DevPF* knockdowns. (A) Nucleotide identity across *DevPF1* (bottom) and *DevPF2* (top) genes. Screenshot of pairwise sequence alignment in Geneious prime software. Silencing region (violet), exon (green), intron (white), perfect identity (gray) and mismatch/gap (black). Scale in base pairs at the top. (B) mRNA expression levels of *DevPF1* (top) and *DevPF2* (bottom) upon KDs (*ND7* (control), *DevPF1* and *DevPF2*) at different developmental time points (onset, early, late and very late). Lines represent the mean of all replicates for a given KD and time point. The cell stage composition of each time point averaged over all KDs is shown at the top (individual compositions in Fig. S5), along with schematic representations of the considered cell stages. (C) Protein expression upon KD: fluorescence intensities of DevPF1-GFP (top) and DevPF2-GFP (bottom). Red line: median. Whiskers: 1.5 times the interquartile range from the lower or upper quartile. Dots: data points outside the whiskers. Sample size = 10. (D) IES retention score (IRS) correlations between *DevPF1*- and *DevPF2*-KD replicates. Diagonal: IRS distributions of individual KDs. Below diagonal: correlation graphs of pairwise comparisons. Above diagonal: corresponding Spearman correlation coefficients. Red lines: ordinary least-squares (OLS) regression, orange lines: LOWESS, and gray lines: orthogonal distance regression (ODR).

The mRNA levels of *DevPF1* and *DevPF2* were examined during a time course experiment (more details and further analysis follow in subsequent sections) (Fig. 4B). Consistent with the published expression profiles (Arnaiz et al., 2017), *DevPF1* expression in the *ND7* control knockdown (KD) was highest during onset of development and gradually declined to almost no expression at the “very late” time point. The late-expressed *DevPF2* peaked at the “late” time point in the control KD. The expression of both genes was strongly reduced upon their respective KDs (*DevPF1* mRNA levels were reduced upon *DevPF1*-KD; *DevPF2* mRNA levels were reduced upon *DevPF2*-KD). A lesser reduction was also observed upon silencing of the respective paralog (*DevPF1* levels were reduced in *DevPF2*-KD and vice versa). Thus, the *DevPF1* and *DevPF2* silencing constructs lead to co-silencing which is less efficient than the target gene silencing.

To investigate how the changes in mRNA levels affect protein levels, we checked the localization of the GFP-tagged DevPFs upon KDs. Since *DevPF1* is expressed throughout the whole development, multiple developmental time points were collected (Fig. S3A). For the late-expressed *DevPF2*, only cell stages with clearly visible new MACs were considered (Fig. S3B). In addition to *ND7*-KD, the knockdown of *PGM*, the gene encoding the PiggyMac IES excisase (Baudry et al., 2009), was performed to test whether the disturbance of IES excision alters DevPF localization. Neither the localization of DevPF1-GFP nor of DevPF2-GFP was impaired by either of the control KDs. In contrast, the GFP signals were almost completely lost upon *DevPF1*- or *DevPF2*-KD. To quantify this observation, GFP fluorescence signals were measured in new MACs (Fig. 4C) as both paralogs exclusively localize to the new MACs during late stages. In line with the observed reduction in mRNA levels, DevPF1-GFP expression was efficiently reduced upon *DevPF1*-KD, whereas *DevPF2*-KD led to a weaker reduction. For DevPF2-GFP, the levels were almost equally reduced in *DevPF1*- and *DevPF2*-KD. Thus, we confirmed co-silencing on both mRNA and protein levels with reduced silencing efficiency compared to the targeted KD. Therefore, all results obtained in KD experiments must be considered, at least in part, as a combined effect of silencing both DevPF1 and DevPF2, albeit with only a partial contribution from the non-targeted gene silencing.

To further investigate the impact of co-silencing on the KD analysis we examined IES retention score (IRS) correlations of multiple KD replicates (more details and further analysis in subsequent sections). The *DevPF2*-KD replicates showed high to moderate correlation among each other while they correlated less well with two out of four *DevPF1*-KD replicates (Fig. 4D). This suggests that despite the partial co-silencing, individual KD effects might be observed.

### *DevPF1* and *DevPF2* affect IES excision genome-wide

The influence of the DevPFs on genome reorganization was initially investigated with survival tests and IES retention PCRs upon KDs. Reduced protein levels during sexual development can induce errors including IES retention, impacting the survival of the subsequent generation. For survival tests, the growth of the cells that completed their sexual development was followed for several divisions. IES retention PCRs test for the presence (failed excision) of specific IESs in the new MAC genome. *ND7*-KD and *PGM*-KD were used as negative and positive control, respectively. To investigate the possibility that the observed effects result from off-target silencing of an unrelated gene, two silencing probes (a and b) were tested for each paralog (Fig. 4A). *DevPF1* and *DevPF2* KDs with either of the silencing probes resembled *PGM*-KD, with high lethality in the new progeny (Fig. 5A) and retention of selected IESs (Fig. 5B). This indicates that both *DevPF1* and *DevPF2* contribute to IES excision.

**Figure 5:**
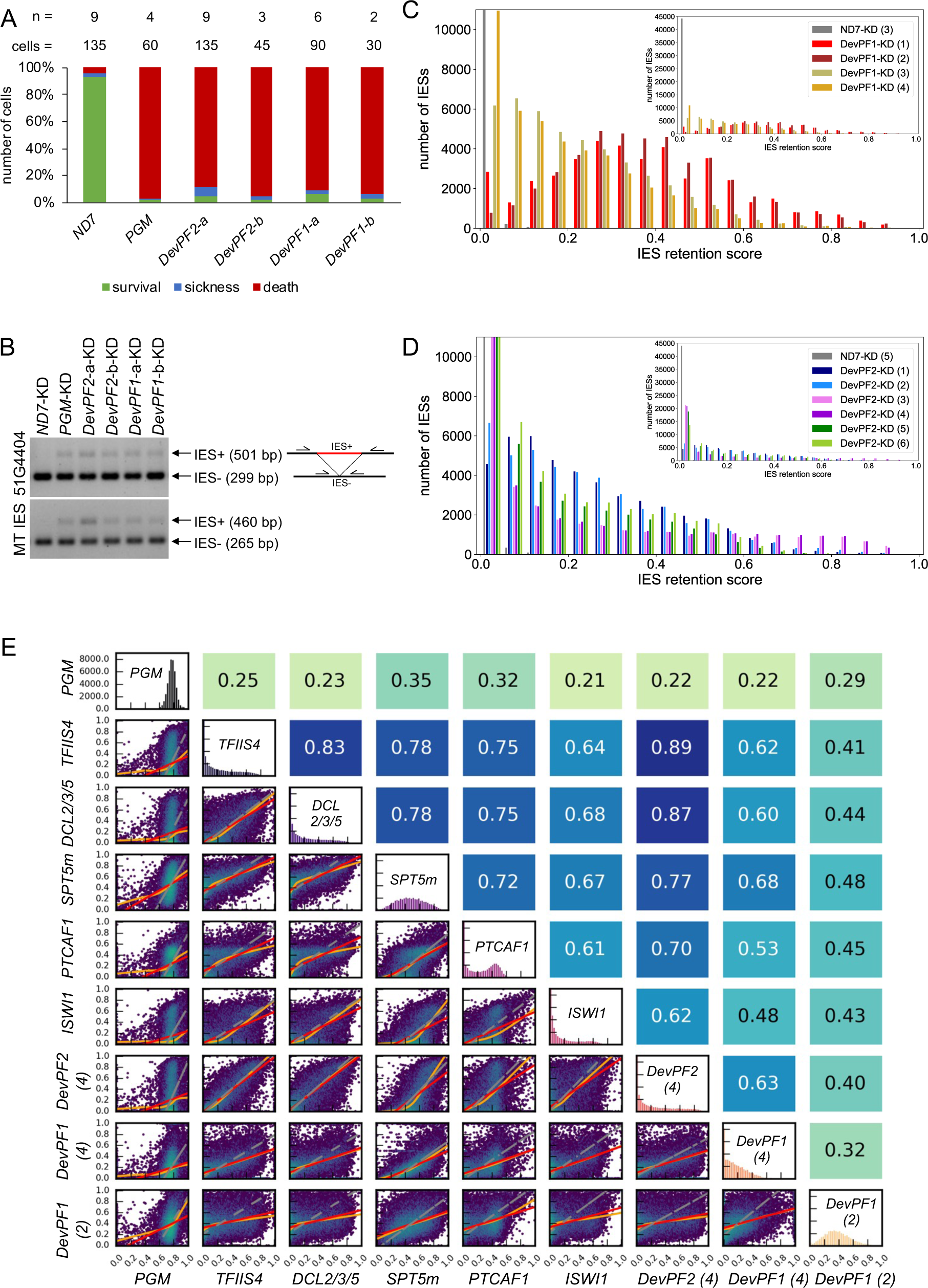
Effects of *DevPF* knockdowns on genome-wide IES retention. (A) Viability of new progeny after KDs (*ND7* (negative control), *PGM* (positive control), *DevPF1* and *DevPF2*) during sexual development. For *DevPF1* and *DevPF2*, two silencing regions were targeted (a and b, see Fig. 4A). The numbers of experiments (n) and cells counted (cells) are indicated at the top. Survival: normal division. Sickness: reduced growth. Death: 3 or less cells after three days. (B) IES retention PCRs for two IESs on genomic DNA isolated from KD cells. (C) and (D): IES retention score (IRS) histograms for *DevPF1* (C) and *DevPF2* (D) KD replicates, indicated in parentheses. (E) IRS correlation between KDs. Diagonal: IRS distributions of individual KDs. Below diagonal: correlation graphs of pairwise comparisons. Above diagonal: corresponding Spearman correlation coefficients. Red lines: ordinary least-squares (OLS) regression, orange lines: LOWESS, and gray lines: orthogonal distance regression (ODR).

Next, we tested genome-wide IES retention in enriched new MAC DNA samples. We observed considerably elevated levels of retained IESs in both *DevPF1*- and *DevPF2*-KD (Fig. 5C,D). Notably, differences between replicates of the same KDs were observed, whereas replicate pairs processed in parallel (see Methods) exhibited similar profiles. Correlations among the paralog replicates indicated that despite varying IES retention distributions, *DevPF2*-KD replicates demonstrated high correlations among themselves (Fig. 4D). *DevPF1*-KD replicates correlated less well with each other, and *DevPF1*-KD replicate 3 (3) showed a high correlation with the *DevPF2*-KDs. This indicates that *DevPF2*-KD replicates were more consistent than the *DevPF1*-KD replicates.

Genes that work closely together are expected to show similar KD effects on IES retention. To identify functionally related genes, *DevPF1*-KD and *DevPF2*-KD IRS data was correlated with published data from other gene KDs (Fig. 5E). *DevPF2*-KD (4) was selected from the *DevPF2* replicates. *DevPF1*-KD (2) and *DevPF1*-KD (4) were selected as representative of the variability observed in the *DevPF1*-KDs. *DevPF2*-KD (4) displayed high correlation with other KDs, such as *TFIIS4* and *DCL2/3/5* (Fig. 5E). Moderate correlation was observed for *DevPF1*-KD (4) with *SPT5m*, whereas *DevPF1*-KD (2) did not correlate well with any of the tested KDs.

Short IESs are proposed to predominantly rely on the excision complex (specifically Pgm (Baudry et al., 2009) and Ku80c (Marmignon et al., 2014)) for removal, while long IESs tend to require additional molecules for excision (Sellis et al., 2021). To determine whether *DevPF1*- and *DevPF2*-KD preferentially affect long IESs, the length distribution of the top 10% of highly retained IESs in each KD was plotted (Fig. S4A,B, Table S1). In comparison to the length distribution of all IESs, *DevPF2*-KD (4) showed an overrepresentation of long IESs, similar to observations in *EZL1*-KD, silencing of the catalytic subunit of the PCR2 complex (Frapporti et al., 2019; Lhuillier-Akakpo et al., 2014), or *DCL2/3/5*-KD, silencing of the scnRNA and iesRNA biogenesis proteins (Lepère et al., 2009; Sandoval et al., 2014), (Fig. S4A). Conversely, the highly retained IESs in *DevPF1*-KD (2) did not show the same overrepresentation and resembled the profile in *PGM*- and *KU80c*-KD, silencing of two members of the excision complex. Again, the replicates of the *DevPF* KDs exhibited variation in the extent of the observed effect (Fig. S4B).

Defects in IES excision not only result in the retention of IESs but can also lead to excision at alternative TA boundaries. So far, alternative excision above background levels has only been reported for silencing of ISWI1 and its complex partners (Singh et al., 2022, 2023). Neither *DevPF1*-KD nor *DevPF2*-KD resulted in elevated levels of alternative excision (Fig. S4; Table S2).

### *DevPF1*- and *DevPF2*-KD alter the small RNA population

The early-produced 25 nt scnRNAs and the late-produced 26-30 nt iesRNAs have been proposed to assist MIC-specific sequence targeting in the new MACs (Sandoval et al., 2014). Therefore, the small RNA populations across developmental time points upon DevPF KDs were analyzed (Figs 6A, S6A). In *DevPF1*-KD (2), scnRNA production was completely abolished, an effect also observed in the KD of genes proposed to be involved in scnRNA production: the two scnRNA-processing genes *DCL2* and *DCL3* (Sandoval et al., 2014), and *STP5m*, involved in the generation of the transcripts serving as substrates for Dcl2/3 cleavage (Gruchota et al., 2017). The KD of the late-expressed *DevPF2* showed a much weaker reduction of scnRNA production, which might be caused by co-silencing of *DevPF1*.

**Figure 6:**
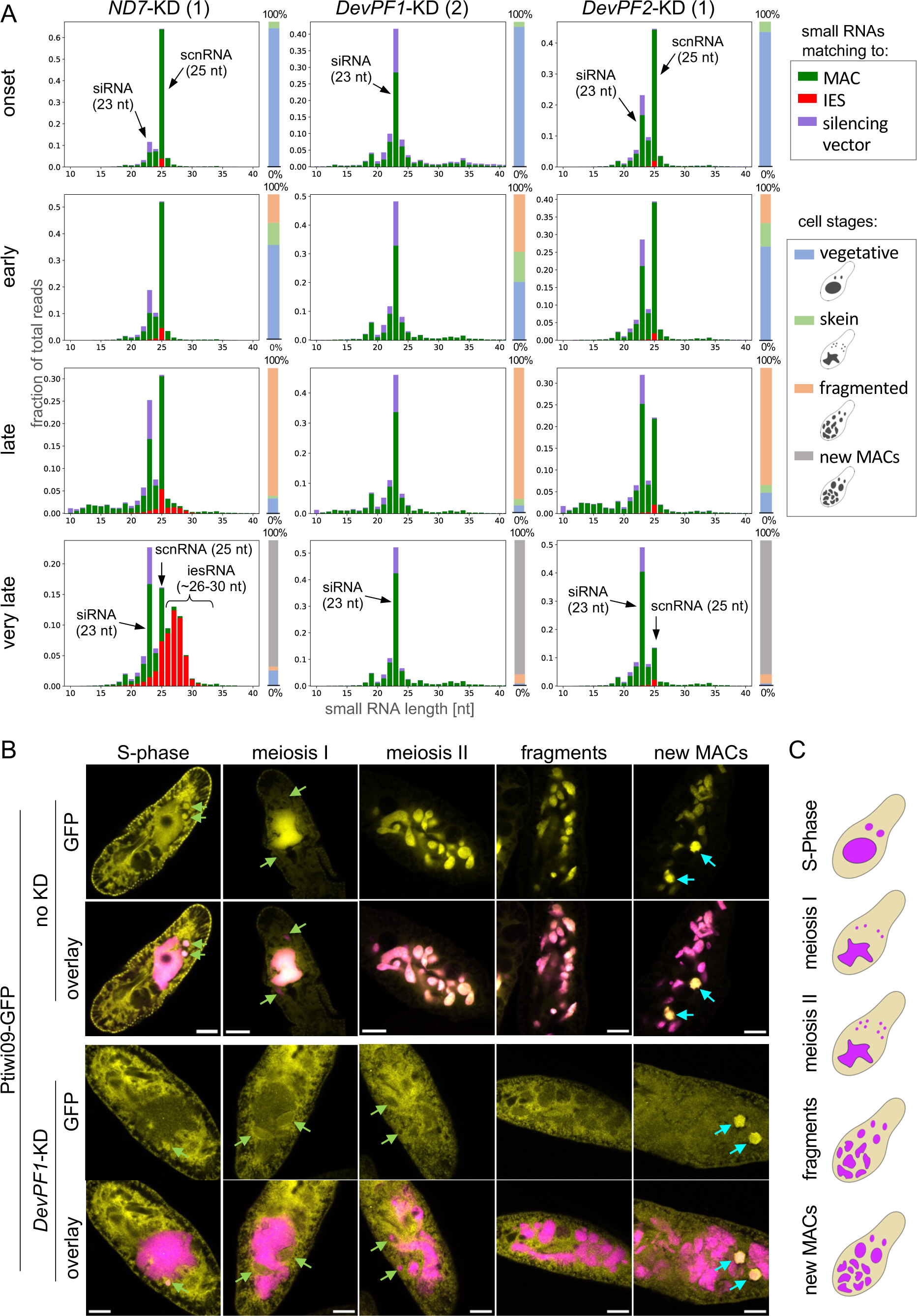
Changes of small RNA populations upon *DevPF* knockdowns. (A) Small RNA populations (10-40 nt) at developmental time points (onset, early, late and very late) in different KDs (*ND7* (control), *DevPF1* and *DevPF2*), mapping to silencing plasmid backbone (vector), MAC or IES sequences. Individual cell stage compositions are indicated by the bar to the right of each diagram, along with schematic representations of the cell stages considered. (B) Ptiwi09-GFP localization at different developmental stages in the context of no (top) and *DevPF1* KD (bottom). DAPI (pink) and GFP (yellow). Individual z-planes for GFP and overlay (DAPI and GFP). Green arrows: MICs. Cyan arrows: new MAC. Scale bar = 10 µm. (C) Schematic representation of cell stages in (B).

To further investigate DevPF1’s effect on the scnRNA pathway, we observed Ptiwi09-GFP localization upon *DevPF1*-KD. Ptiwi09, together with Ptiwi01, stabilizes the scnRNAs throughout scnRNA selection in the parental MAC and targeting of MIC-specific sequences in the new MACs (Bouhouche et al., 2011; Furrer et al., 2017). As previously described (Bouhouche et al., 2011; Singh et al., 2023), Ptiwi09-GFP localizes first to the cytoplasm and parental MAC with a transient localization in the swelling MICs before shifting to the new MAC (Fig. 6B). Upon *DevPF1*-KD (Fig. 6B), the localization to the MICs before meiosis I is not impaired; however, the translocation into the parental MAC is strongly reduced and Ptiwi09-GFP predominantly remains in the cytoplasm throughout meiosis II and MAC fragmentation. We have reported a similar change in Ptiwi09-GFP localization upon *DCL2/3*-KD (Singh et al., 2023), suggesting that the loss of scnRNAs is responsible for the failed protein transfer into the parental MAC. Similar to *DCL2/3*-KD, *DevPF1* depletion does not affect Ptiwi09-GFP’s localization to the new MACs (Fig. 6B).

Interestingly, DevPF1-HA IP at two developmental time points (early: about 30% fragmentation; late: visible new MACs in fragmented cells) identified Ptiwi01/09 as potential interaction partners of DevPF1 with a higher enrichment in the early than the late time point (Fig. S6B, Table S3). None of the other small RNA-related proteins were detected (Dcls, Spt5m, TFIIS4 or Ptiwi10/11).

For both *DevPF1-* and *DevPF2*-KD, iesRNA production was impaired. iesRNAs are proposed to derive from dsRNAs transcribed from excised IESs (Allen et al., 2017; Sandoval et al., 2014). Hence, failed excision of IESs in *DevPF1*- or *DevPF2*-KD contributes to reduced iesRNA levels, as has consistently been observed for many other KDs of genes involved in *Paramecium* genome editing (Charmant et al., 2023; de Vanssay et al., 2020; Ignarski et al., 2014; Maliszewska-Olejniczak et al., 2015; Singh et al., 2022; Wang et al., 2023). The lack of scnRNAs in the *DevPF1*-KD cannot explain the absence of iesRNAs, as these accumulate even if the preceding scnRNA production is blocked (Sandoval et al., 2014). In the late time point analyzed for DevPF IPs, peptides mapping to Ptiwi10/11/06 were detected in DevPF2-IP (Fig. S6C, Table S3), but not DevPF1-IP (Fig. S6B). Therefore, DevPF2 might contribute to iesRNA biogenesis by an interaction with Ptiwi proteins.

### *DevPF1*- and *DevPF2*-KD affect mRNA expression

Since PHD fingers have often been reported to be involved in gene expression regulation (Aasland et al., 1995; Sanchez & Zhou, 2011) we sought to investigate whether the *DevPF* KDs alter mRNA expression levels during development. Batch effects had a major influence on the variance within the replicates (Fig. S7A), as observed for IES retention (Fig. 5C,D).

*DevPF1*-KD showed almost no differentially expressed genes compared to *ND7*-KD during onset of development (Fig. 7A). During this early stage, genes are transcribed solely from the parental MAC, where DevPF1-GFP does not localize (Figs 2A, 3). Surprisingly, in *DevPF2*-KD, a high number of genes were differentially expressed during the onset of development (Fig. 7A). Since *DevPF2* is late-expressed and *DevPF1*-KD showed no effect, the observed difference might be caused by differing cell stages within the collected populations of *DevPF2*-KD and *ND7*-KD. During the “early”, “late” and “very late” time points, *DevPF1*- and *DevPF2*-KD showed similar changes in mRNA expression.

**Figure 7:**
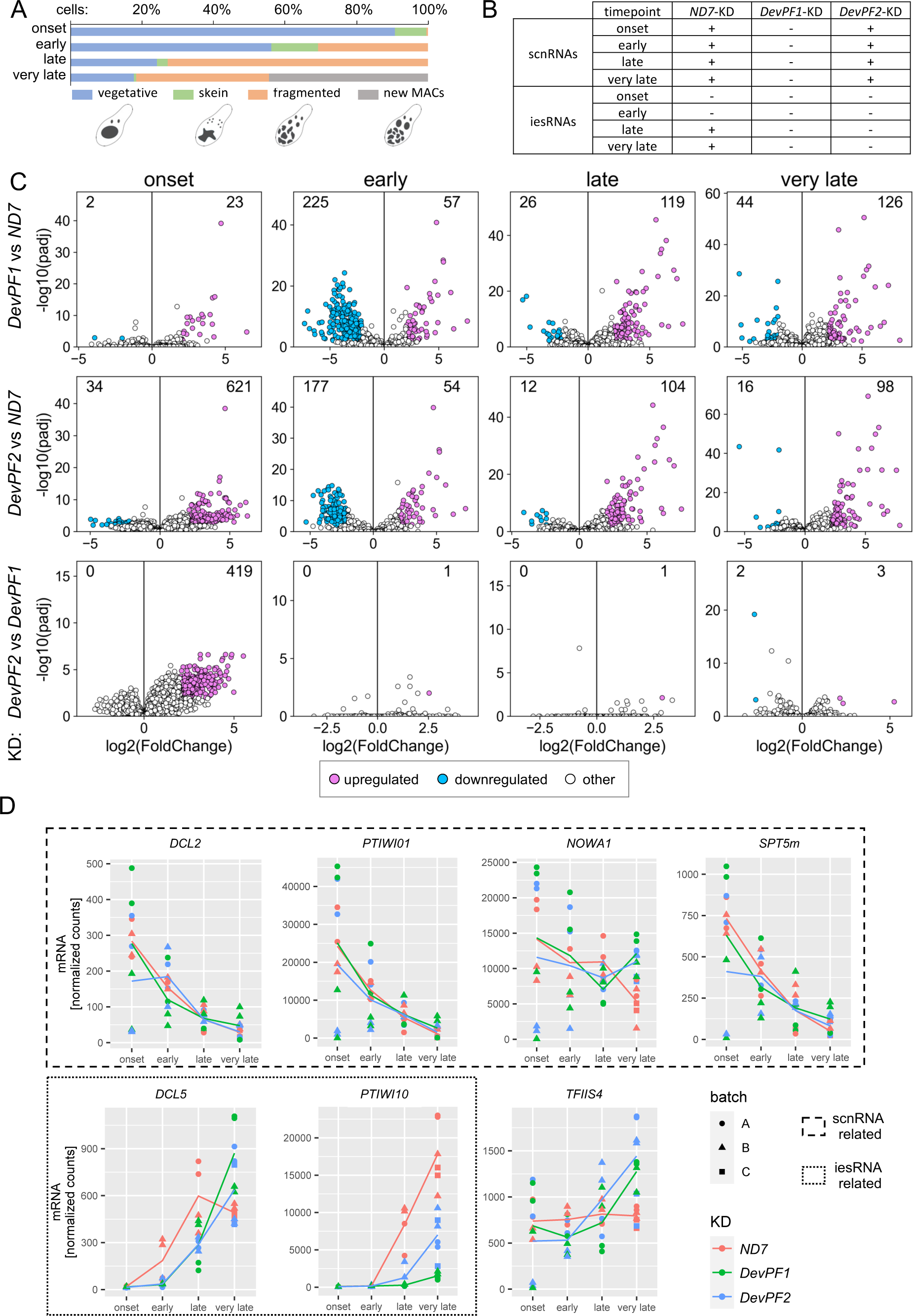
Differential gene expression in *DevPF* knockdowns. (A) Cell stage composition of each time point averaged over all KDs (individual compositions in Fig. S5), along with schematic representations of the considered cell stages. (B) Presence or absence of scnRNAs and iesRNAs in different KDs (*ND7*, *DevPF1* and *DevPF2*) and time points (onset, early, late, very late). (C) Differentially expressed genes in *DevPF1*-(top) or *DevPF2*-(middle) compared to *ND7*-KD or *DevPF1*-compared to *DevPF2*-KD (bottom) at different developmental time points (onset, early, late and very late). Thresholds for up-/downregulation: adjusted p-value < 0.01; |log2(fold change)| > 2. The number of up-/downregulated genes is indicated in each diagram. For all comparisons, 35777 transcripts were analyzed, except for: DevPF1-ND7 onset (33696), DevPF2-ND7 early (35083), and DevPF2-DevPF1 onset (34389). (D) Gene expression levels of selected genes upon KDs (*ND7* (control), *DevPF1* and *DevPF2*) at different developmental time points (onset, early, late and very late). The lines represent the mean of all replicates in a given KD and time point.

The abolishment of development-specific small RNAs in the *DevPF*-KDs might result from downregulation of genes involved in scnRNA or iesRNA production. We observed no general trend indicating a drastic reduction of expression of scnRNA-related genes, like *DCL2*, *PTIWI01* or *SPT5m* (Figs 7B, S7B, Table S4, S5). However, these trends in expression should be considered with the caveat of considerable expression variability and limitation of the number of replicates that could practically be obtained. At least for Ptiwi09, the localization experiments upon *DevPF1*-KD confirmed no loss in protein levels (Fig. 6B).

The expression of iesRNA-related genes was altered in both *DevPF1*- and *DevPF2*-KD compared to *ND7*-KD (Figs 7D, S7B, Table S4, S5). *DCL5*, the Dicer-like protein responsible for the initial cleavage of IES derived dsRNAs into small iesRNAs (Sandoval et al., 2014), was downregulated (Table S4, S5) in early stages, but tended to be upregulated in the very late stage (Table S4, S5). *PTIWI10* and *PTIWI11*, the Piwi proteins responsible for further processing and stabilization of iesRNAs during the positive feedback loop (Furrer et al., 2017), were downregulated in both *DevPF1*- and *DevPF2*-KD (Table S4, S5). Successful expression of *PTIWI10/11* has been proposed to depend on IES excision since both genes are expressed from the new MAC and harbor IESs in their flanking/coding regions (Furrer et al., 2017) (Fig. S7C). If IES retention was the only cause for downregulation, one would expect higher IRSs for these IESs in KDs with lower mRNA levels. While the mRNA reduction is stronger in *DevPF1*-KD than in *DevPF2*-KD (Fig. 7D, Table S4, S5), this trend is not reflected in the IRSs of the IESs whose retention is proposed to interfere with *PTIWI10/11* expression (Table 1). In most of the KD replicates, there is no or low retention (IRS < 0.1) and the replicates showing moderate to high retention (0.1 < IRS < 0.3) belong to both *DevPF1*-and *DevPF2*-KD. Hence, the reduced mRNA levels of *PTIWI10/11* cannot only be explained by IES retention.

**Table 1:**
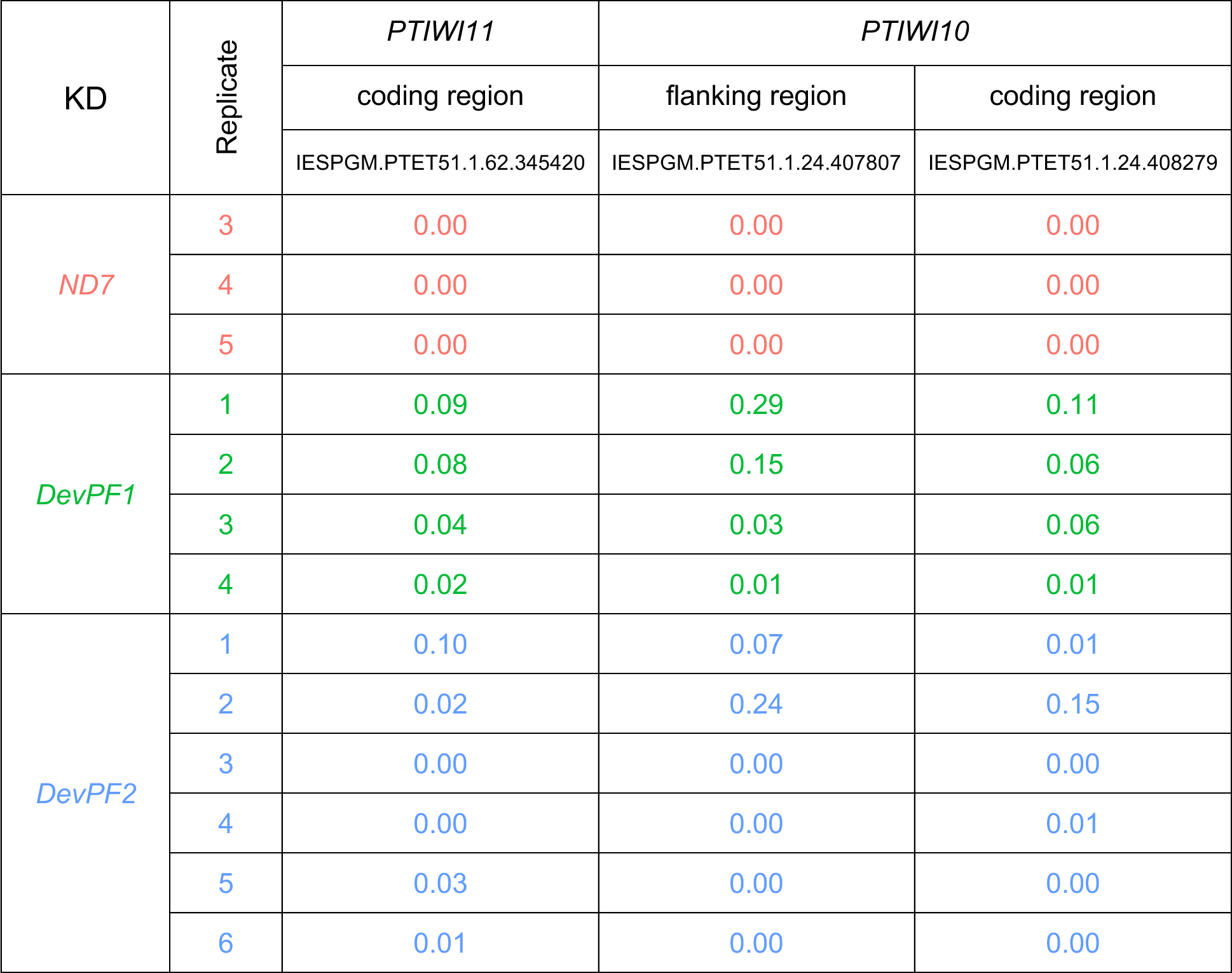
IES retention scores of IESs at *PTIWI10/11* genes. The genes *PTIWI10* and *PTIWI11* contain IESs in their coding and/or flanking regions, which were proposed to impair their transcription when retained. The IRS values for the three relevant IESs (IDs with prefix IESPGM.PTET51.1) are provided for each KD. Rows are color-coded according to the KDs as shown in the mRNA read count diagrams (i. e. Figs 7D, S7B).

## Discussion

### Implications of the PHD domain for DevPF1 and DevPF2 functions

Genome reorganization is a fundamental process underlying cell and immune system development and some diseases (Bassing et al., 2002; Forment et al., 2012; Mani & Chinnaiyan, 2010; Rooney et al., 2004). Ciliates undergo massive genome reorganization during the maturation of their somatic genome. This makes them excellent models for studying the complex mechanisms involved in the targeted elimination of genomic sequences (Beisson et al., 2010d). In the present study, we described two paralogous PHD finger proteins, DevPF1 and DevPF2, involved in IES excision in *Paramecium*. Both paralogs harbor a PHD and a PHD zinc finger-like domain (Fig. 1). These domains belong to the zinc-finger family and the PHD domain is characterized by a well-conserved C4HC3 motif (Aasland et al., 1995; Schindler et al., 1993). The eight core amino acids of this motif coordinate two zinc ions and thereby provide structural stability to the domain (Pascual et al., 2000). Among other histone-binding domains, such as bromodomains or PWWP domains, PHD fingers are the smallest (Fleck et al., 2021; Miller et al., 1985). Multiple sequence alignment and structure predictions confirmed the presence of the characteristic C4HC3 motif in both DevPF1 and DevPF2 (Fig. 1), suggesting that both PHDs might be functional.

PHD fingers, mainly nuclear proteins, are often considered epigenetic readers, recognizing histone modifications, primarily on the histone 3 (H3) N-terminal tail (Sanchez & Zhou, 2011). Peptides matching to histones were enriched in the DevPF-IPs of late developmental time points (Fig. S6B,D, Table S3), however none of them was specific to H3. PHD fingers have been reported to bind non-H3 partners, like DNA, histone 4, or other proteins (Bienz, 2006; Black & Kutateladze, 2023; Gaurav & Kutateladze, 2023; L. Liu et al., 2012; Oppikofer et al., 2017). The combination of the PHD and PHD-zinc-finger-like domain in the DevPFs may enable the paralogs to simultaneously recognize two adjacent histone modifications, as demonstrated for tandem PHD domains (Zeng et al., 2010). PHD domains are also found in various chromatin associated proteins involved in gene regulation. Notably, ISWI-containing chromatin remodeling complexes often include a subunit with a PHD domain, such as the ACF (Eberharter et al., 2004), NURF (Haitao Li et al., 2006; Wysocka et al., 2006) or WICH (Bozhenok et al., 2002) complexes.

DevPF2 was initially identified in pulldowns of the ISWI1 protein, and, thus far, no PHD-containing protein has been shown to be a part of this remodeling complex (Singh et al., 2022, 2023). It is intriguing to consider that DevPF2 might contribute PHD functionality to the ISWI1 chromatin remodeling complex. However, *DevPF2*-KD does not show elevated levels of alternative excision (Fig. S4C-E) that are characteristic of other members of the complex so far (Singh et al., 2022, 2023) and ISWI1 was not identified as a potential interaction partner in the DevPF2-IP (Fig. S6C). If DevPF2 interacts with the ISWI1 complex, we infer that it may not be a core complex component, particularly as it does not contribute to excision precision.

### A potential role for DevPF1 and DevPF2 as transcription factors?

#### A potential role in non-coding transcription in the MICs (for scnRNA production)

DevPF1’s localization in the MICs (Figs 2A, 3) and its importance for scnRNA production (Fig. 6A) could point towards its involvement in the bidirectional transcription of the MIC genome for scnRNA production. Spt5m (Gruchota et al., 2017) and TFIIS2/3 (Maliszewska-Olejniczak et al., 2015) are proposed to be involved in this micronuclear transcription. One of the *DevPF1*-KD replicates showed moderate IRS correlation with *SPT5m* (Fig. 5E) (to our knowledge, no IRS data exists for TFIIS2 or TFIIS3) and *SPT5m*-KD also reduces scnRNA production. The localization of Dcl2-GFP (Lepère et al., 2009), Ptiwi09-GFP (Fig. 6B) and DevPF1-GFP (Fig. 2A) in the swelling MICs suggests that scnRNA biogenesis occurs during the S-phase of meiosis. Ptiwi09 and DevPF1 may interact in the MICs or the cytoplasm. Non-crosslinked IP’s would be needed to further verify this interaction. However, *PTIWI01/09*-KD does not completely abolish scnRNAs (Furrer et al., 2017), indicating that DevPF1 acts upstream of scnRNA loading and guide strand removal. Future investigations of bi-directional transcription and scnRNA biogenesis will allow to identify how all these molecules cooperate.

Spt5m-GFP, TFIIS2/3-GFP and DevPF1-GFP are present in the MICs beyond S-phase and localize to the new MACs at later stages (Gruchota et al., 2017; Maliszewska-Olejniczak et al., 2015). Their role in the MIC during meiotic divisions remains unknown. It was speculated that Spt5m might be involved in co-transcriptional deposition of epigenetic marks that sustain meiotic processes, ultimately aiding in IES targeting. The potential of PHD domains to bind histone modifications raises a similar possibility for DevPF1. However, its role appears to be more specific, as DevPF1 is not present in all gametic and zygotic nuclei simultaneously (Fig. 2&3).

Msh4/5, homologs of proteins essential for crossover, are also present in all gametic nuclei during the first and second meiotic division, and their silencing leads to substantial IES retention (Rzeszutek et al., 2022). However, their non-canonical functions that lead to IES retention are not yet fully understood (Rzeszutek et al., 2022). Since new MACs develop in *DevPF1*-KD (Figs 6B,S3) and *MSH5*-KD cells, neither of the genes are essential for crossover or karyogamy. More research will be needed in future to decipher the functions of the DevPF proteins in the gametic nuclei.

#### A potential role in non-coding transcription in the new MAC (for scnRNA-based targeting and iesRNA production)

Non-coding transcription in the new MAC, which is hypothized to generate substrates for scnRNA pairing, was proposed to be regulated by the putative transcription elongation factor TFIIS4 that specifically localizes to the early new MACs (Maliszewska-Olejniczak et al., 2015). *DevPF2*-KD IRSs of some replicates correlated most strongly with *TFIIS4*-KD (Fig. 5E), pointing towards a shared functionality. Both DevPF1 and DevPF2 have the potential to act in the same regulatory process as TFIIS4 because both their GFP-fusions localize to the new MACs. In fact, there are reports of transcription factors that combine the TFIIS and PHD domains: Bypass of Ess1 (Bye1) protein in S*accharomyces cerevisiae* harbors a PHD and a TFIIS-like domain, with the former recognizing histone 3 lysine 4 trimethylation and the latter establishing contact with polymerase II for transcriptional regulation (Kinkelin et al., 2013; Pinskaya et al., 2014). It is possible that similar functionality is separated on two individual proteins in *Paramecium*. However, TFIIS4 was not detected in either of the DevPF-IPs in the late developmental stage.

The production of iesRNAs was also proposed to depend on the non-coding transcription of concatenated excised IES fragments (Allen et al., 2017; Sandoval et al., 2014). Although it was established that IES concatemers are likely formed by DNA ligase 4 (Lig4) (Allen et al., 2017), little is known about the proposed bidirectional transcription to produce substrates for Dcl5 cleavage. Allen *et al*. speculated on the involvement of TFIIS4. Since iesRNA production is almost completely abolished in *DevPF1*- and *DevPF2*-KD, a contribution to this transcription is plausible.

The potential function of the DevPFs may extend far beyond TFIIS4-dependent transcription: whereas TFIIS4-GFP localizes transiently to early new MACs (Maliszewska-Olejniczak et al., 2015), DevPF2-GFP and DevPF1-GFP remain in the new MACs for much longer (Fig. 2).

### A potential role in gene transcription in the parental and the new MAC

Early in development, the parental MAC is solely responsible for gene expression and, after genome reorganization progresses, the new MAC contributes at later stages (Berger, 1973). In *Tetrahymena*, E2F family transcription factors were shown to control the cell cycle through gene expression during meiosis (Zhang et al., 2018). DevPF1 and DevPF2 are unlikely to be active in the parental MAC since none of the GFP-fusion proteins localized there (Fig. 2). Consistently, *DevPF1*-KD showed no differential gene expression compared to *ND7*-KD during the onset of development (Fig. 7C) and Ptiwi09-GFP expression was not impaired upon *DevPF1*-KD (Fig. 6B). However, it is difficult to reach a definite conclusion for other genes due to the high variability in expression between the replicates (Figs 7D, S7B) and the high number of differentially expressed genes in *DevPF2*-KD (Fig. 7C) observed during the onset of development. Cells in the “onset” time point are challenging to collect because cell staging relies on MAC morphology changes visualized by DAPI staining. Truly vegetative cells cannot be distinguished from cells initiating meiosis since their MACs look the same; however, the gene expression profiles are expected to differ substantially (Figs 2A, S2B). The collection of subsequent time points is more reliable because the alteration of old MAC shape as development progresses is pronounced.

At the subsequent stages, *DevPF1*- and *DevPF2*-KD affected similar genes. Either, the changes are nonspecific to the *DevPF*-KDs and result from the proposed nuclear crosstalk to adjust transcription levels to accommodate for failed IES excision (Bazin-Gélis et al., 2023) or they are specific to the *DevPF*-KDs and both paralogs exhibit similar functions in the regulation of gene expression. Interestingly, differential expression was observed at the “early” time point (Fig. 7C). *GTSF1*-KD, also causing substantial IES retention, hardly shows any differentially expressed genes at a comparable stage (*DevPF1/2*-KD: 282/231 differentially expressed genes, respectively, at about 30% fragmentation (Fig. 7C); *GTSF1*-KD: 10 differentially expressed genes at about 30-50% fragmentation; (Wang et al., 2023)). This indicates that the early change in gene expression might be specific to *DevPF*-KDs, potentially mediated by other proteins shuttling into the parental MAC. Since Ptiwi09-GFP translocates efficiently to the parental MAC upon *GTSF1*-KD (Wang et al., 2023) but not upon *DevPF1*-KD (Fig. 6B), it might be worth investigating differential expression upon *PTIWI01/09*-KD.

Late in development, gene expression starts from the new MACs (Berger, 1973), where both DevPF paralogs localized (Fig. 2). Some late-expressed genes, like *PTIWI10*, are expressed only from the new MAC after the initial onset of IES excision (Furrer et al., 2017). Indeed, *PTIWI10/11* mRNA levels are downregulated in *DevPF1*-KD or *DevPF2*-KD (Fig. 7D, S7B, Table S4, S5). This trend cannot be explained solely by the strength of retention observed for the IESs interfering with *PTIWI10/11* expression (Table 1). It suggests that DevPF1 and DevPF2 may regulate gene expression in the new MAC, albeit specifically for some genes like *PTIWI10* and *PTIWI11.* The extent of gene expression regulation by the DevPFs beyond these genes remains uncertain. To further investigate if the DevPFs serve as transcription factors, and if so, which genes they regulate, genes associated with DevPF binding could be identified by techniques like Cut-and-Run (Skene et al., 2018) and compared to mRNA expression changes upon *DevPF*-KDs.

### Potential cytoplasmic functions

In contrast to the other putative transcription factors discussed so far (Spt5m, TFIIS2/3/4, DevPF2), DevPF1-GFP exhibits a pronounced cytoplasmic distribution in the early stages of development (Fig. 2A). While most described PHD fingers are nuclear proteins, some can be recruited to the cytoplasm or plasma membrane by binding partners (Betz et al., 2004; Gozani et al., 2003). DevPF1 may play a role in transmitting signals of sensed starvation to the MICs, initiating sexual development. As *DevPF1* is not constitutively expressed during vegetative growth (Figs 1B, S2B), another factor is needed to first initiate *DevPF1*’s gene expression in the parental MAC. However, DevPF1 might interact with specific markers of starvation in the cytosol, promoting early sexual processes. If that is the case, DevPF1 is not essential for general meiotic processes, as meiosis and new MACs development show no defects in *DevPF1*-depleted cells (Figs 6B,S3). Since peptides matching Ptiwi01/09 were identified in the DevPF1-IP, the Ptiwi01/09 complex is a potential binding partner of DevPF1 in the cytoplasm. However, since Ptiwi01/09 are highly expressed proteins (Bouhouche et al., 2011), further IP experiments would be needed to verify this interaction.

### DevPF1’s selective localization to gametic and post-zygotic nuclei

The selective localization of DevPF1 to certain gametic and post-zygotic nuclei (Fig. 3) raises intriguing questions about its potential role in nuclear fate decisions. The survival and destruction of the gametic nuclei depends on their subcellular positioning (Grandchamp & Beisson, 1981). DevPF1 may play a role in either promoting their movement or preparing for their degradation. However, the observed number of nuclei simultaneously containing DevPF1-GFP (zero to four) neither fits the number of nuclei selected for survival (one) nor for degradation (seven). DevPF1 may either contribute to this process successively or may not be directly related to the nuclear fate itself. The fate of the post-zygotic nuclei is decided during the second mitotic division by the subcellular localization of the division products (Grandchamp & Beisson, 1981). This means, from each post-zygotic nucleus, one of the division products will remain as MIC and one develops into a new MAC. During the second mitotic division, DevPF1-GFP was observed in one of the two dividing nuclei. Its localization in the precursor of one MIC and one MAC without being present in the precursor of the other MIC and MAC, does not imply its involvement in the nuclear fate decision. Furthermore, *DevPF1*-KD neither impaired the selection of gametic nuclei nor the differentiation of the new MACs.

The specific localization of nuclear proteins to certain nuclei in multinuclear cells has been studied extensively in insect embryos. In *Drosophila*, the transcription factors Bicoid (Driever & Nüsslein-Volhard, 1988) and Dorsal (Roth et al., 1989) establish the anterior-posterior, and dorsal-ventral axis, respectively, by initiating gene expression depending on the cytoplasmic localization of the nuclei. The activity of the transcription factors is restricted by gradients to a certain cytoplasmic region (Morisato & Anderson, 1995; Spirov et al., 2009). However, DevPF1-GFP’s nuclear localization does not appear associated with subcellular localization of the nuclei and it remains unclear how DevPF1-GFP is specifically recruited.

As only fixed cells were examined, the dynamics of DevPF1-GFP localization were not captured. The fact that DevPF1-GFP localization is independent of nuclear divisions (Fig. 3B), combined with observations of cells at the meiotic or mitotic division stage with an absence of DevPF1-GFP in all nuclei (Fig. S2C), suggests that DevPF1 localization might be asynchronous and transient. Possibly it is recruited to each of the gametic nuclei at some point before the completion of the second meiotic division and to each of the post-zygotic nuclei before completion of the second mitotic divisions. Live cell imaging could illuminate the dynamics of DevPF1 localization and its correlation with nuclear fate. However, this approach presents challenges, as it requires confocal imaging to capture the DevPF1-GFP signal in the MICs, and the observation time scale would need to span across multiple hours of *Paramecium* development.

### DevPF1: a general factor for IES excision

DevPF1 plays a role throughout sexual development: from the early stages before meiosis to the very late stages (Fig. 2A). It appears to influence various aspects of genome reorganization in the MICs and the new MACs, including scnRNA production and potentially expression of certain genes. Consequently, the depletion of *DevPF1* affects the excision of a wide range of IESs (Fig. 5C). However, it is important to reiterate that we observed high batch-to-batch variability in the *DevPF* replicates in both IES retention (Fig. 5C, D) and mRNA expression (Figs 7D, S7B). The time point collection had a major influence on mRNA levels (Fig. S7A). Variable new MAC enrichment by a sucrose gradient might introduce variation into the IRS analysis, as fragments of the parental MAC add unexcised IES sequences, diluting the effect of IES retention (Charmant et al., 2023). Fluorescence-activated nuclear sorting (FANS) enables better nuclear separation in *Paramecium* (Charmant et al., 2023; Guérin et al., 2017) and should be able to eliminate most of such variation. Additionally, microinjection of DNA into macronuclei before RNAi experiments can be used to control for contaminating DNA from old MAC fragments.

Revisiting previous KD experiments with additional replicates would be worthwhile to explore the extent of batch-to-batch IRS and expression variance for other KDs. Noteworthy, variability in IES retention across replicates has recently been shown for *GTSF1* (Charmant et al., 2023; Wang et al., 2023), suggesting this phenomenon is not restricted to *DevPF1* and *DevPF2*. In general, KD experiments are challenging to tightly control for reproducibility, and more effort should be invested in generating knockouts in *Paramecium*, as established in *Tetrahymena* (Chalker, 2012).

It has been shown that evolutionarily old IESs tend to be short and are excised early in development, independent of additional factors apart from the excision machinery (Sellis et al., 2021). On the other hand, evolutionarily young IESs tend to be long, later excised and dependent on the scnRNA pathway and the deposition of histone modifications in the new MAC for their excision (Sellis et al., 2021; Swart et al., 2014; Zangarelli et al., 2022). In line with this, most gene KDs tested in this study exhibited an overrepresentation of long IESs among their most highly retained IESs, including *DevPF2* (Fig. S4A,B). Only *PGM*-KD, *KU80c*-KD and two of the *DevPF1*-KD replicates showed no preference for long IESs. Pgm and Ku80c are components of the excision machinery and are therefore expected to affect all IESs. While DevPF1 may not be a direct part of the excision machinery, it appears to have a general contribution to IES excision, regardless of the length of the IES. Consequently, we propose that DevPF2 contributes to the excision of long IESs, while DevPF1 may serve as a more general factor.

## Methods

### Paramecium tetraurelia cultivation

Mating type 7 (MT7) cells from strain 51 of *Paramecium tetraurelia* were grown in Wheat Grass Powder (WGP, Pines International) medium supplemented with 10 mM sodium phosphate buffer (pH 7.3). WGP medium was bacterized with *E.coli* strain HT115 to feed paramecia, and the cultures were maintained either at 27°C or at 18°C according to the standard protocol (Beisson et al., 2010b, 2010c).

### Protein localization imaging by fluorescence microscopy

Plasmids for microinjection were generated by amplifying the coding and flanking sequences from MT7 genomic DNA and introducing them with the PCR-based method CPEC (Quan & Tian, 2011) into the L4440 plasmid (Addgene, USA). DevPF1 was expressed with its endogenous flanking regions (304 bp upstream of the DevPF1 start codon and 272 bp downstream of the DevPF1 stop codon). DevPF2 endogenous flanking regions (455 bp upstream the DevPF2 start codon and 273 bp downstream of the DevPF2 stop codon) yielded no expression. Therefore, as *PGM* exhibits a similar expression profile to *DevPF2* (Fig. 1B), DevPF2 genomic coding sequence was inserted between the *PGM* flanking regions (96 bp upstream of the *PGM* start codon and 54 bp downstream of the *PGM* stop codon). Before the stop codon, the *GFP* coding sequence was connected to the protein coding sequences via a glycine-serine-linker (SSGGGSGGSGGGS). 60 μg of plasmid DNA was linearized with AhdI (New England Biolabs, UK) and extracted with phenol-chloroform for injection.

*Paramecia* were microinjected with either C-terminally GFP-tagged DevPF1 (endogenous regulatory regions) or C-terminally GFP-tagged DevPF2 (*PGM* regulatory regions) following the standard protocol (Beisson et al., 2010a). Sexual development was induced by starvation and cells of different developmental stages were collected and stored in 70% ethanol at −20°C. To stain cells with DAPI (4,6-diamidino-2-2-phenylindole), cells were dried on a microscopy slide, washed twice with phosphate-buffered saline (PBS) and permeabilized for 10 min at RT (room temperature) with 1% Triton X-100 in PHEM (PIPES, HEPES, EGTA, magnesium sulfate), fixed with 2% paraformaldehyde (PFA) in PHEM and washed once for 5 min at (RT) with 3% BSA (bovine serum albumin, Merck-Sigma, Germany) in Tris-buffered saline with 10 mM EGTA and 2 mM MgCl_2_ (TBSTEM). After DAPI (2 μg/ml in 3% BSA) incubation for 7-10 min at RT, the cells were mounted 40 µl of ProLong Gold Antifade mounting medium (Invitrogen, USA) or ProLong Glass Antifade mounting medium (Invitrogen, USA). For α-tubulin staining, after permeabilization and fixation, cells were blocked for 1 h at RT with 3% BSA and 0.1% Triton X-100. Primary rat anti-α-tubulin antibody (Abcam, UK) was diluted 1:200 in 3% BSA and 0.1% Triton X-100 in TBSTEM and incubated overnight at 4°C. After 3 washes with 3% BSA, the goat anti-rat secondary antibody conjugated to Alexa fluorophore 568 (Abcam, UK) was diluted 1:500 in 3% BSA and 0.1% Triton X-100 in TBSTEM and incubated for 1 h at RT. After two washes, cells were stained with DAPI and mounted with Prolong Glass Antifade mounting medium.

Images were acquired on a confocal SP8 Leica fluorescence microscope (60x/1.4 oil objective) with constant laser settings. The detector (photon multiplier) gain for the DAPI signal (430-470 nm) varied to accommodate differences in signal strength (500-550 V). Postprocessing was done in Fiji (version 2.14.0/1.54f) (Schindelin et al., 2012). Brightness and contrast in the GFP channel was set the same in all the images to be compared (Figs 2, S2B: DevPF1-GFP: Min 0, Max 681 and DevPF2-GFP: Min 0, Max 170; Figs 3, S2C: DevPF1-GFP: Min 0, Max 703; Fig. S3: constant settings for each cell stage).

### Knockdown efficiency validation using fluorescence intensity

Cells injected with either DevPF2-GFP or DevPF1-GFP were subjected to KDs of *ND7*, *PGM*, *DevPF2* and *DevPF1* genes. Cells during new MAC development were collected (for details see methods on silencing experiments), then stained with DAPI and mounted on ProLong Glass Antifade as described above. Images of a single z-plane through the new MAC were acquired on a SP8 Leica Confocal microscope with 60x/1.4 oil objective using the same laser settings for all images. For each KD, 10 cells were imaged. In Fiji software (version 2.14.0/1.54f), the brightness and contrast in the GFP channel was set the same values for all images compared in the same analysis (DevPF1-GFP injected cells: Min 0, Max 1078; DevPF2-GFP injected cells: Min 0, Max 298). Fluorescent signal was measured in a constant area in 1 MAC of each cell and the area mean was used as intensity for this nucleus. The area was set in the DAPI channel and the fluorescence was measured in the GFP channel. Since the same area was measured for each nucleus, no normalization was used to account for nuclear size variation. To account for background fluorescence, GFP fluorescence in non-transformed wild type cells was measured and the mean of all wild type cells was subtracted from all measured intensities. All intensities were normalized to the mean of all *ND7*-KD cells in the corresponding injection. All scripts are available from https://github.com/Swart-lab/DevPF_code.

### Co-immunoprecipitation

Paramecia were injected with either Human influenza hemagglutinin (HA)-tagged DevPF1 (same cloning strategy as described before) or GFP-tagged DevPF2. For DevPF1-HA, an early time point (about 30% fragmentation) and late time point (new MACs clearly visible in fragmented cells) was collected, while for DevPF2-GFP, only the late time point was collected. Non-transformed wild type cells were collected as controls. Cells were washed twice with 10 mM Tris and as much liquid was removed as possible. For 300 ml initial culture volume, cells were fixed with 1 ml 1% PFA for 10 min at RT and quenched with 100 µl of 1.25 M glycine for 5 min at RT. After one wash with PBS (centrifugation for 1 min at 4°C and 1000 g), 2 ml lysis buffer (50 mM Tris, 150 mM NaCl, 5 mM MgCl_2_, 1% Triton X-100, 10% Glycerol and cOmplete protease inhibitor EDTA-free (Roche, Germany)) were added and cells were sonicated using an MS72 tip on a Bandelin Sonopulse device with 52% amplitude for 15 s on ice. The pellet and input fraction were separated by centrifugation (13,000 g, 4°C, 30 min).

To enrich HA-tagged proteins, 50 µl beads (Anti-HA-affinity matrix, Merck-Sigma, Germany) were washed thrice (500 g, 4°C, 2 min) in ice-cold IP buffer (10 mM Tris pH 8, 150 mM NaCl, 1 mM MgCl_2_, 0.01% NP-40, 5% Glycerol, cOmplete protease inhibitor EDTA-free (Roche, Germany) and incubated with 1 ml of cleared input lysate overnight at 4°C. After four washes with ice-cold IP buffer, the bound proteins were eluted from the beads in 50 µl 2× PLB (10% SDS, 0.25 M Tris pH 6.8, 50% Glycerol, 0.2 M DTT, 0.25% Bromophenol blue) at 98°C for 20 min (IP fraction).

To enrich GFP-tagged proteins, 25 µl beads (GFP-Trap Agarose beads, Chromotek, Germany) were washed once with ice-cold 20 mM Tris pH 7.5 with 100mM NaCl (2,500 g, 4°C, 5 min) and thrice in ice-cold IP buffer. Beads were incubated with 1 ml cleared input lysate for 1 to 2 h at 4°C and washed four times with ice-cold IP buffer. Bound proteins were eluted in 30 µl 2× PLB at 98°C for 20 min (IP fraction).

For western blots, 0.5% of total input and 15% of total IP fraction were resolved on 10% SDS-PAGE gels and wet transferred onto a 0.45 µm nitrocellulose membrane for 2 h at 80 V and 4°C (Bio-Rad, Germany). The membrane was blocked for 1 h in 5% BSA in PBST (PBS + 0.2% Tween20). HA-tagged proteins were detected with an HRP-conjugated anti-HA antibody (sc-7392 HRP, Santa Cruz, USA) diluted 1:500 in PBST and incubated overnight at 4°C. GFP-tagged proteins were detected with an primary anti-GFP antibody (ab290, Abcam, UK) diluted 1:2000 and incubated overnight at 4°C followed by an secondary anti-rabbit HRP conjugated antibody (12-348, Merck Millipore, Germany) diluted 1:5000 in PBST and incubated for 1 h at RT. Membranes were screened using AI600 (GE Healthcare, Germany).

Samples were sent to EMBL’s Proteomics Core Facility in Germany for mass spectrometry experiments and analysis. Using R, contaminants were removed from the FragPipe output files (protein.tsv, (Kong et al., 2017)), and only proteins quantified with a minimum of two razor peptides were included for subsequent analysis. After log2 transformation of raw TMT reporter ion intensities, batch effect correction (limma package’s (Ritchie et al., 2015) ‘removeBatchEffects’ function), and variance stabilization normalization (vsn) with vsn package (Huber et al., 2002), the abundance difference in WT and DevPF samples was maintained by determining different normalization coefficients. To investigate differential protein expression (limma package), replicate information was incorporated in the design matrix with the ‘lmFit’ limma function. “hit” annotation: false discovery rate (FDR) smaller 5% and a fold change of at least 100%. “candidate” annotation: FDR smaller 20% and a fold change of at least 50%. Scripts to generate volcano plots are available from https://github.com/Swart-lab/DevPF_code.

### Silencing experiments, survival test and IES retention PCR

Silencing constructs for *DevPF2* and *DevPF1* were generated by cloning genomic gene fragments into a T444T plasmid (Sturm et al., 2018) (Addgene, USA) using CPEC (Quan & Tian, 2011). For both *DevPF1* and *DevPF2*, two silencing regions were selected: DevPF1 silencing region a (525 bp fragment from 3-527; position 1 is the first nucleotide of the start codon); DevPF1 silencing region b (733 bp fragment from 532-1264); DevPF2 silencing region a (525 bp fragment from 3-527); DevPF2 silencing region b (731 bp fragment from 532-1262). Co-silencing was predicted with the RNAi off-target tool from ParameciumDB (Heng Li & Durbin, 2009) for both silencing regions (*DevPF1* silencing region a and b: 19 and 30 hits, respectively, in *DevPF2* gene; *DevPF2* silencing region a and b: 19 and 30 hits, respectively, in *DevPF1* gene). The plasmids were transformed into HT1115 (DE3) *E. coli* strain and expression was induced overnight at 30°C with Isopropyl ß-D-1-thiogalactopyranoside (IPTG; Carl Roth, Germany). Paramecia were seeded into the silencing medium at a density of 100 cells/ml to induce sexual development by starvation after 4 to 6 divisions. KD experiments were performed as previously described (Beisson et al., 2010e).

After the paramecia finished sexual development, 15 cells were transferred into a regular, non-induced, feeding medium for the survival test. Paramecia were monitored for three days to observe growth effects. For IES retention PCRs, genomic DNA was extracted from cultures that finished sexual development using GeneElute – Mammalian Genomic DNA Miniprep Kit (Merck-Sigma, Germany). PCRs were done on specific genomic regions flanking an IES (Table S6) to check for the retention of IESs. 1-12.5 ng DNA was used as input andPCR products were resolved on 1-2% agarose gels.

### Time course silencing experiments

The time course experiments were conducted in three batches, each processing two KD replicates in parallel (batch A: replicates 1 and 2 of *ND7*-, *DevPF1*- and *DevPF2*-KD; batch B: replicates 3 and 4 of *ND7*-, *DevPF1*- and *DevPF2*-KD; batch C: replicates 5 and 6 of *ND7*- and *DevPF2*-KD). In batch A and B, cells were collected as soon as the first meiotic cells were observed in the population (onset), between 20 to 40% fragmentation (early), at 80-90% fragmentation (late) and 6 h after the late time point (very late). In batch C, cells were collected before the onset of autogamy (vegetative), at 50% fragmentation (early), at 100% fragmentation + visible anlagen (very late) and 6 h later (very late + 6h). Since batch C was collected at different stages, only the “very late” time point of batch C was considered for differential expression analysis. For all time course replicates, enriched new MAC DNA was analyzed for IES retention and total RNA was collected from the collected time points for sRNA and/or mRNA analysis.

### Macronuclear isolation and Illumina DNA-sequencing

Samples for new MAC isolation were collected from the KD cultures of all time course experiments three days after completion of sexual development as described previously (Arnaiz et al., 2012). DNA library preparation (350 bp fragment sizes) and Illumina sequencing (paired-end, 150 bp reads) were done at Novogene (UK) Company Limited, Cambridge according to their standard protocols.

### IES retention and alternative boundary analysis

For IES retention score analysis, whole genome sequencing reads of enriched new MAC DNA after KD were adaptor trimmed using TrimGalore (Krueger, 2019) if significant Illumina adapter content was observed using FastQC v0.11.9 (Andrews, 2010) (see Table S7 for adapter sequences). The “Map” module of ParTIES v1.05 pipeline was used to map the reads on MAC and MAC+IES reference genomes with changes in the /lib/PARTIES/Map.pm file as described in (Singh et al., 2023). The IES retention scores (IRS) were calculated by the “MIRET” module (provided as DevPF_IRS.tab.gz). All scripts are available from https://github.com/Swart-lab/DevPF_code. IRS correlations were calculated as described previously (Swart et al., 2014).

Alternative excision was analyzed as described previously (Singh et al., 2023). In brief, properly paired and mapped reads were selected from the output from the ParTIES “Map” module for the MAC+IES reference genome and downsampled to the same library size (DevPF1-KD (1) and DevPF2-KD (2) were excluded due to small library size). We then employed the “MILORD” module of a pre-release version of ParTIES (13 August 2015) with default parameters to annotate alternative and cryptic IES excision. All scripts are available from https://github.com/Swart-lab/DevPF_code.

The data generate in this study was compared with data of previously published KDs: *PGM*-KD (Arnaiz et al., 2012), *TFIIS4*-KD (Maliszewska-Olejniczak et al., 2015), *SPT5m*-KD (Gruchota et al., 2017), *PTCAF1*-KD (Ignarski et al., 2014), *DCL2/3/5*-KD (Sandoval et al., 2014), *KU80c*-KD (Abello et al., 2020), *EZL1*-KD (Lhuillier-Akakpo et al., 2014) and *ISWI1*-KD (Singh et al., 2022).

### RNA extraction and sequencing

Total RNA was either extracted with phenol-chloroform followed by Monarch Total RNA Miniprep kit (New England Biolabs) or with the Quick-RNA Miniprep kit (Zymo). For phenol-chloroform extraction (batch C), 300 ml cells subjected to RNAi were washed twice with 10 mM Tris pH 7.5 (RT, 280 g, 2 min) and shock frozen by dropping them directly into liquid nitrogen. 500 μl of 2× DNA/RNA protection reagent from the Monarch kit were added to the frozen pellet and the cells thawed by vortexing. After adding 10 μl proteinase K and 1 ml RNA lysis buffer, the manufacturer’s instructions (RNA Binding and Elution (Cultured Mammalian Cells)) were followed. On-column DNase I treatment was included.

For RNA extraction with Quick-RNA Miniprep kit (batch A and B), 100 ml of *Paramecium* cultures subjected to RNAi by feeding were washed twice in 10 mM Tris pH 7.5 in pear-shaped oil flasks by centrifugation (RT, 280 g, 2 min). After the final wash, cells were collected on ice and spun at 2,000 g for 2 min and 4°C and as much liquid as possible was removed. 3× volume of 1× DNA/RNA Shield (Biozym) was mixed with the cells and the samples were stored at −70°C until further processing. For RNA extraction, samples were thawed at RT and mixed with 1× volume of RNA lysis buffer. The manufacturer’s instructions were followed (section: (III) Total RNA Purification). Extracted total RNA was send to Azenta Life Sciences for library preparation (sRNA: NEBNext Small RNA Library Prep Set for Illumina; mRNA: NEBNext Ultra II RNA Library Prep Kit for Illumina) and paired-end Illumina sequencing (NovaSeq 2×150bp).

### Small RNA analysis

Small RNA sequencing reads were trimmed using cutadapt (Martin, 2011) version 3.2 with the parameter -a “AGATCGGAAGAGCACACGTCTGAACTCCAGTCA” to remove the relevant Illumina adaptor sequence. Trimmed reads were mapped to the *Paramecium tetraurelia* strain 51 MAC + IES genome and L4440 (*ND7*-KD) or T444T (*DevPF1*/*DevPF2*-KD) silencing vector with bwa version 0.7.17-r1188 (Heng Li & Durbin, 2009). GNU grep (version 2.14) was used to select 10-49 bp long, uniquely mapped reads (possessing the SAM file format flags “XT:A:U”) and sRNA length histograms were generated by a Python script. All scripts are available from https://github.com/Swart-lab/DevPF_code.

### mRNA analysis

Illumina adapter sequences (Table S7) were trimmed from reads with TrimGalore (Krueger, 2019). Reads were mapped to the *Paramecium tetraurelia* strain 51 transcriptome with hisat2 (Kim et al., 2019) allowing 20 multimappings (-k 20). Using samtools (Heng Li et al., 2009), the properly paired and mapped reads were filtered (-f2 flag) and sorted by the read name (-n flag). Unique mapping reads were acquired with eXpress (Roberts & Pachter, 2013) with 5 additional online expectation-maximization rounds to perform on the data after the initial online round (-O 5 flag) to improve accuracy. Scripts are available from https://github.com/Swart-lab/DevPF_code.

Read counts were normalized with DEseq2 (Love et al., 2014) package in R (version 3.6.3). For plotting, DEseq2 in-build functions plotPCA, plotMA and plotCounts were combined with ggplot2 (Villanueva & Chen, 2019) package (version 3.4.3). Differentially expressed genes were identified for each time point with a Wald test (false discovery rate (alpha) = 0.1). Differentially expressed genes were filtered with an absolute log2(Fold Change) > 2 (corresponding to a 4-fold change) and an adjusted p-value < 0.01. The time point, KD and batch were known sources of variation in the dataset (design = ∼ batch + timepoint + KD+ timepoint:KD). All scripts are available from https://github.com/Swart-lab/DevPF_code.

### Structure prediction with AlphaFold

Protein structures were predicted with AlphaFold2 multimer (Evans et al., 2021; Jumper et al., 2021) using the ColabFold v1.5.2-patch (Mirdita et al., 2022) in Google Colab with default parameters.

### Sequence alignment

Domains were predicted using InterProScan (Paysan-Lafosse et al., 2023). The nucleotide sequence of DevPF2 and DevPF1 (including introns) were aligned with clustalOmega (Sievers et al., 2011) (version 1.2.3) pairwise sequence alignment tool in Geneious prime (version 2023.2.1) with default parameters (Fig 4A).

Multiple sequence alignment of PHD domains was done with clustalOmega (version 1.2.1) using the MPI bioinformatics toolkit’s web interface (Zimmermann et al., 2018) with default parameters.

### Manuscript writing

Grammar and language refinement were assisted by an AI language model developed by OpenAI (GPT-3.5 architecture) (OpenAI, 2023).

## Acknowledgements

We thank the BioOptics core facility and Genome center of MPI for Biology (Tübingen, Germany) for their assistance and Andre Noll for computer system administration.

## Data availability

Supplementary files, including uncropped blot images, microcopy raw files and IES retention scores have been deposited to the open research data repository of the Max Planck Society EDMOND (https://doi.org/10.17617/3.VKJBJ0). Sequencing raw files have been deposited to the European Nucleotide Archive (ENA; https://www.ebi.ac.uk/ena/browser/home) (Leinonen et al., 2011) (accession number: PRJEB67678). The mass spectrometry proteomics data have been deposited to the ProteomeXchange Consortium (Deutsch et al., 2023) via the PRIDE (Perez-Riverol et al., 2022) partner repository (accession number: PXD046704).

