## Supplementary Files for "Two paralogous PHD finger proteins participate in *Paramecium tetraurelia*’s natural genome editing"

### Supplementary information

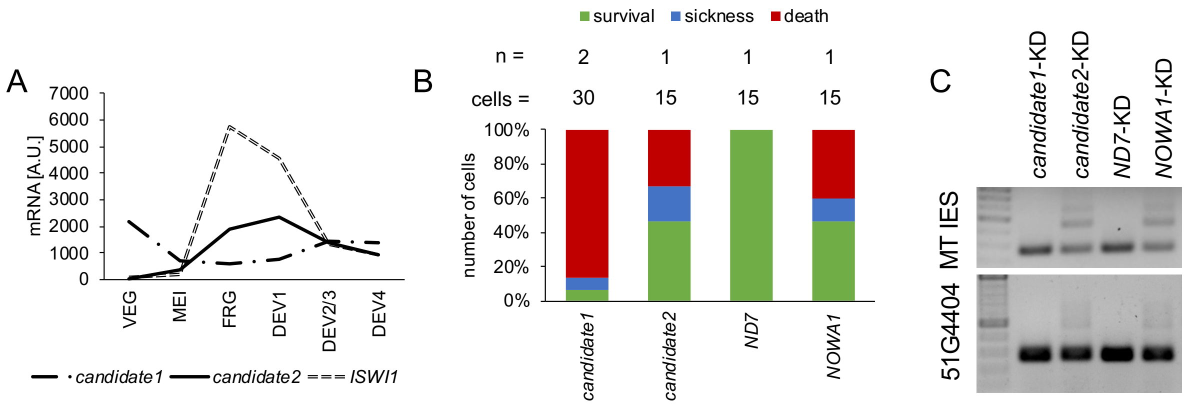


**Figure S1: Screening for new candidate genes involved in *Paramecium* genome reorganization**

(A) mRNA expression profiles in arbitrary units for candidate 1, candidate 2 and *ISWI1* during various developmental stages: VEG (vegetative growth), MEI (micronuclear meiosis and macronuclear fragmentation), FRG (~50% of the population with fragmented maternal MACs), DEV1 (significant proportion with visible anlagen), DEV2/3 (majority with visible anlagen), DEV4 (majority with visible anlagen). Expression data retrieved from ParameciumDB (Arnaiz *et al.*, 2017). (B) Viability of new progeny after knockdowns (*ND7* (negative control) and *NOWA1* (positive control) compared to two candidate genes). Survival: no growth defects. Sickness: reduced division rate. Death: 3 or less cells after three days. The numbers of experiments (n) and cells counted (cells) are indicated at the top. (C) IES retention PCRs for two IESs on genomic DNA isolated from knockdown cells.

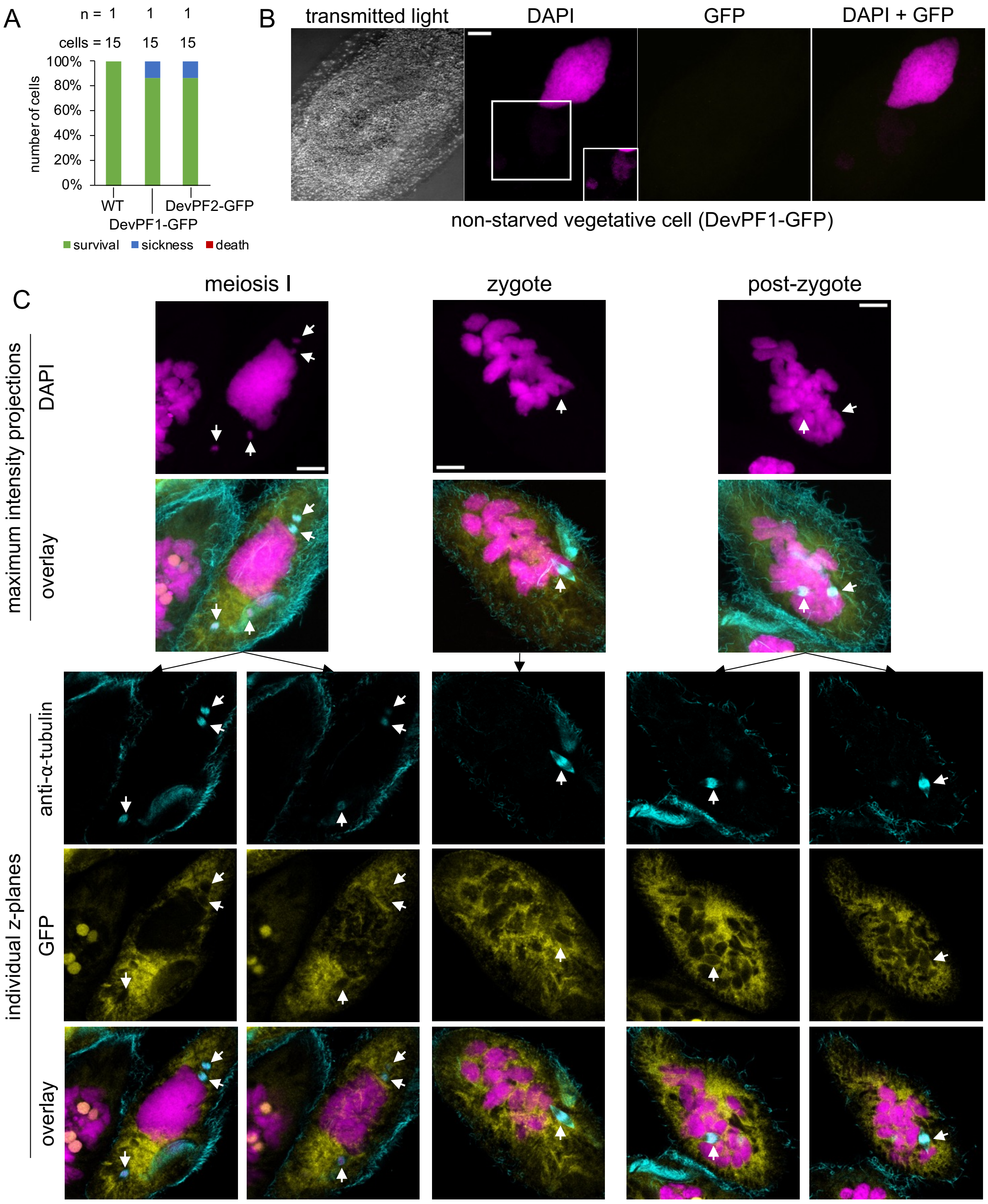


**Figure S2: Additional imaging of DevPF1-GFP localization**

(A) Viability of cells injected with GFP-fusion proteins compared to non-transformed wild type (WT). Survival: normal division. Sickness: reduced growth. Death: 3 or less cells after three days. The numbers of experiments (n) and cells counted (cells) are indicated at the top. (B) DevPF1-GFP localization in non-starved vegetative cells with bacteria inside food vacuoles. (C) Gametic and post-zygotic nuclei lacking DevPF1-GFP localization. Maximum intensity projections of multiple confocal zplanes for (B) and (C) and individual confocal planes for (C). (B) and (C): DNA (stained with DAPI) in pink. GFP in yellow. Scale bar = 10 µm.

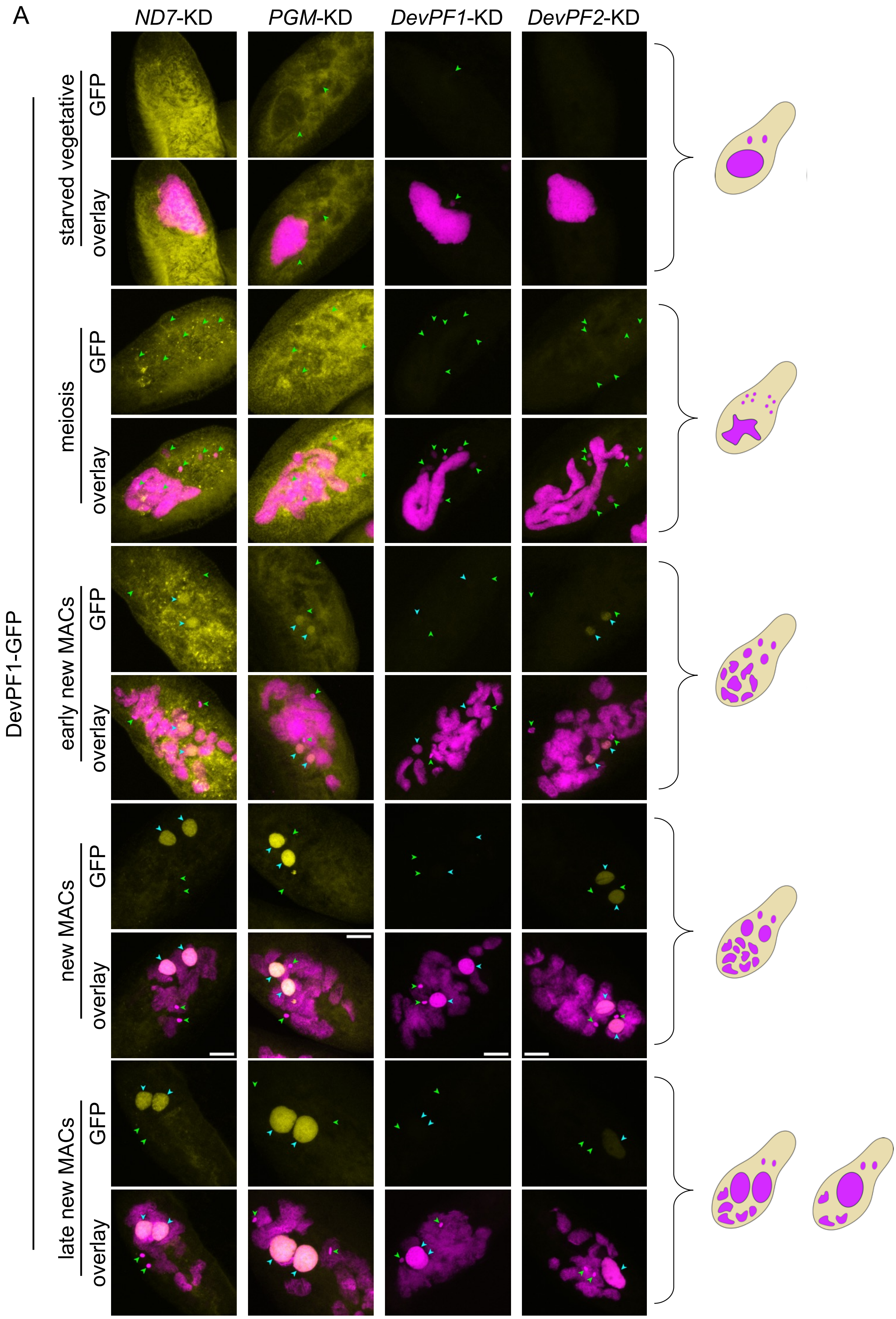

A
ND7-KD
PGM-KD
DevPF1-KD
DevPF2-KD
DevPF1-GFP
starved vegetative
meiosis
early new MACs
new MACs
late new MACs
GFP
overlay

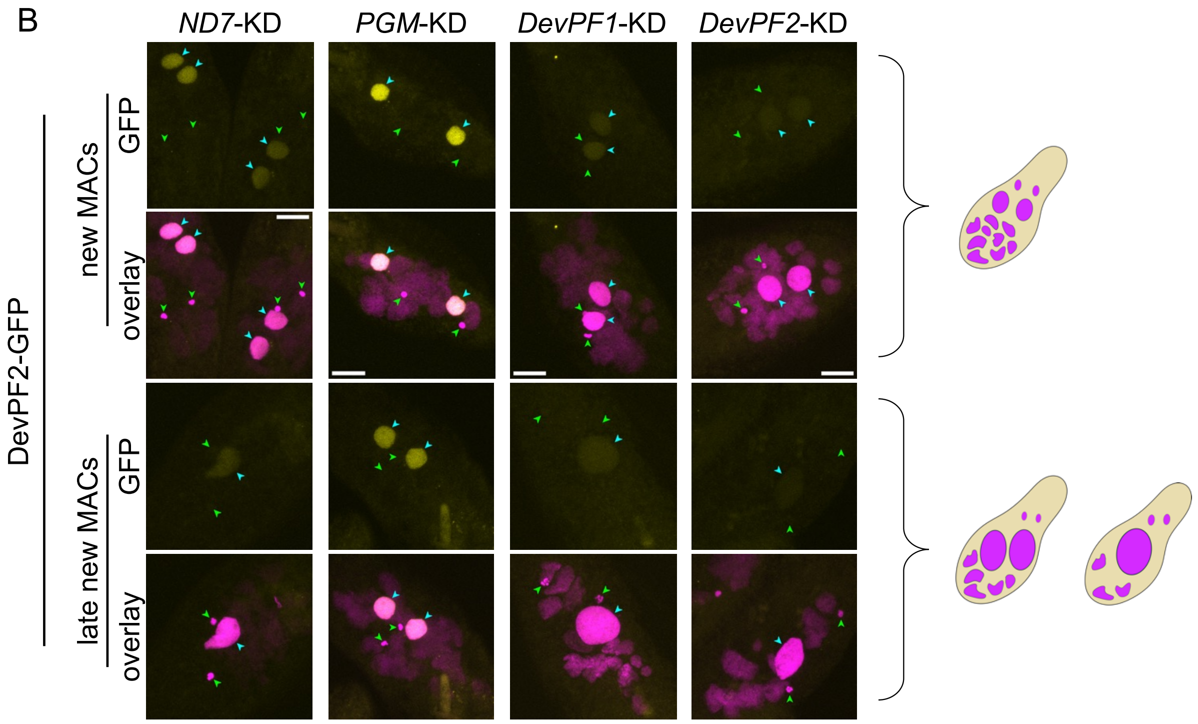


**Figure S3: Localization of DevPF-GFP proteins upon gene knockdowns**

DevPF1-GFP (A) and DevPF2-GFP (B) localization upon *ND7*, *PGM*, *DevPF1* and *DevPF2* knockdown. GFP signal (yellow) and DNA stained with DAPI (pink). Scale bar = 10 µm. Schematic representation of the corresponding cell stage (right).

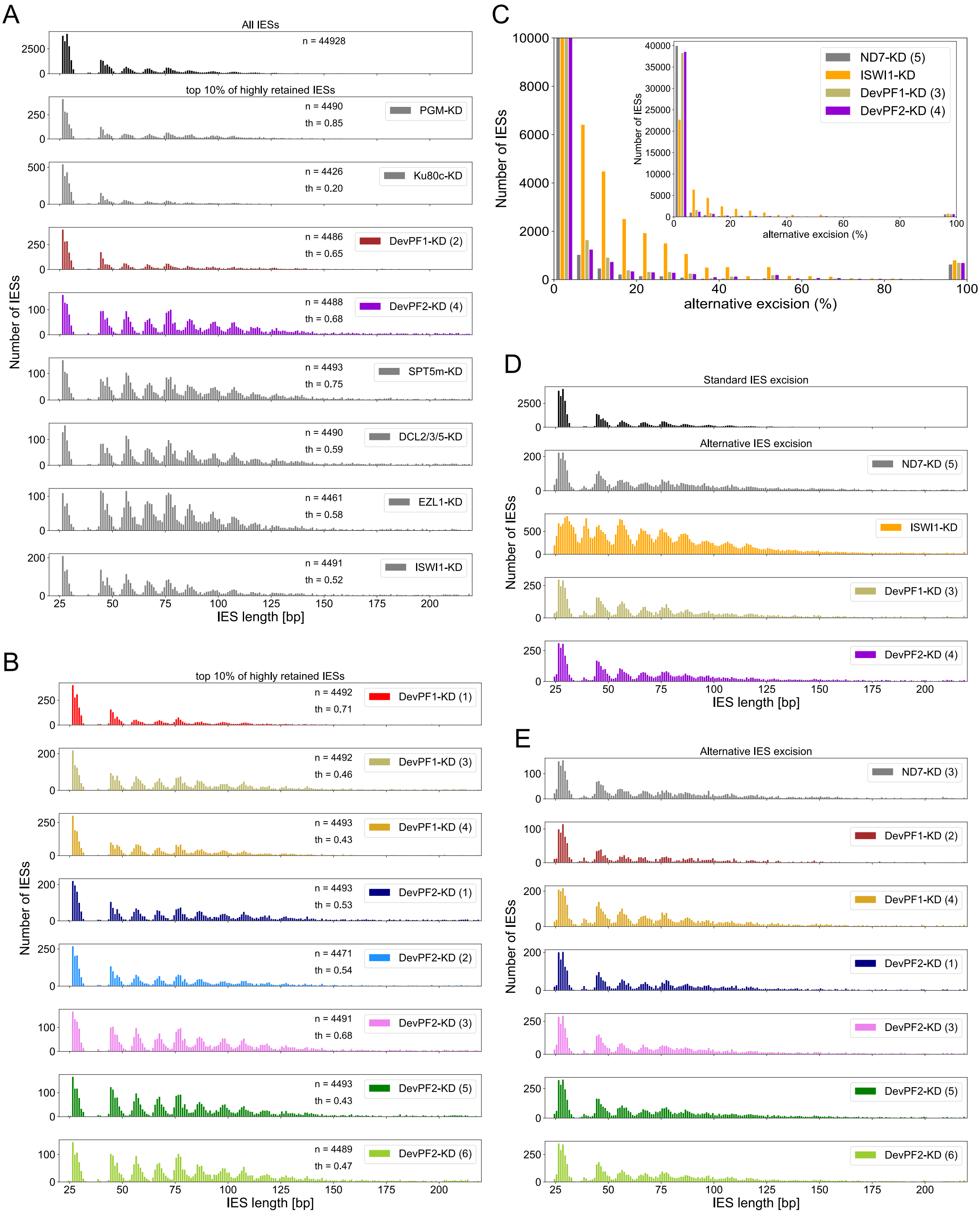


**Figure S4: Analysis of length distributions of standard and alternatively excised IESs upon gene knockdown**

(A) and (B): Length distribution of the top 10% most highly retained IESs in different knockdowns (A) and *DevPF* replicates (B). Indicated are the number of IESs (n) and the IRS threshold (th). (C) Histogram showing the fraction of alternative excision (%) for each IES in various knockdowns. *ND7:* negative control, *ISWI1*: positive control.

(D) and (E): Length distribution of alternatively excised IES in different knockdowns (D) and *DevPF* replicates (E).

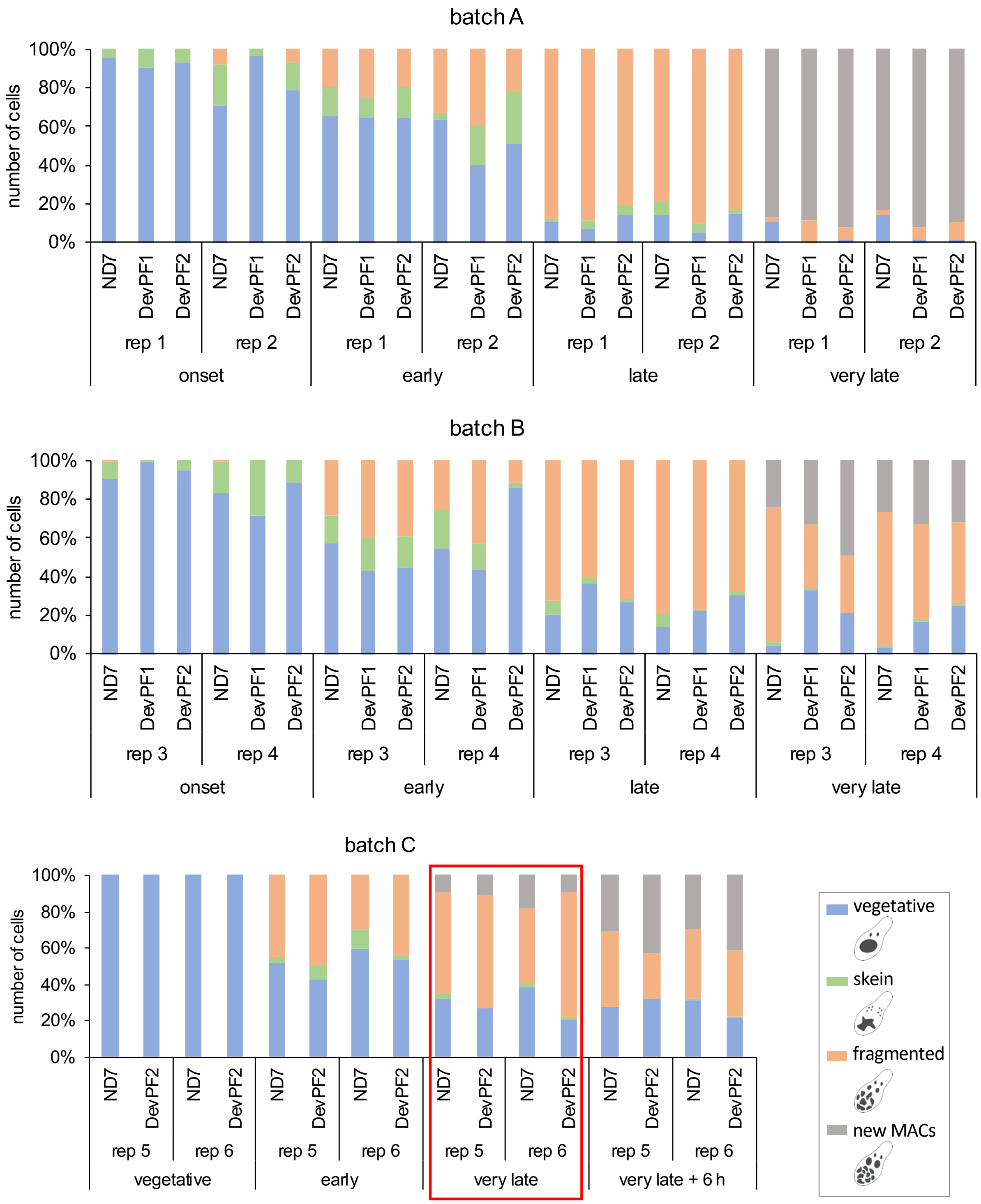


**Figure S5: Cell staging for gene knockdown time course experiments**

Cell staging of 6 time series conducted in 3 batches. A and B included *ND7*-KD, *DevPF1*-KD and *DevPF2*-KD, while C included *ND7*-KD and *DevPF2*-KD. All replicates and time points of batch A and B and “very late” time point of batch C were used for mRNA differential expression analysis. Cells with visible new MACs were considered in the “very late” and “very late + 6 h” time points. The schematic representation of the considered cell stages is provided.

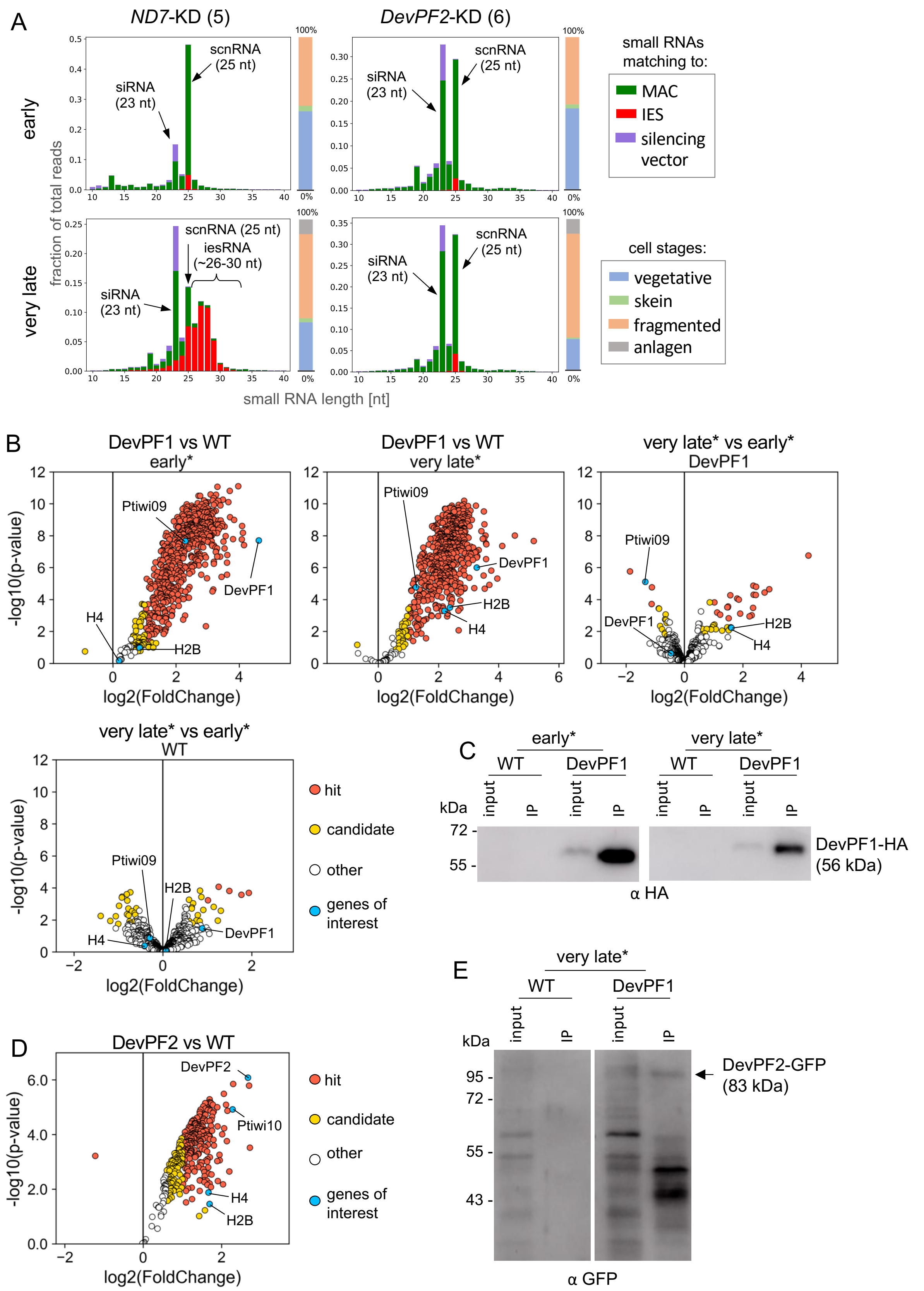


**Figure S6: Small RNA populations and potential DevPF interaction partners**

(A) Small RNA populations (10-40 nt) at “early” and “very late” time points in different knockdowns (*ND7* (control) and *DevPF2*) mapping to silencing plasmid backbone (vector), MAC or IES sequences. Individual cell stage compositions are indicated in the bar to the right of each diagram, along with schematic representations of the cell stages considered. (B) and (D): Genes identified by mass spectrometry in DevPF1-HA (B) or DevPF2-GFP (D) immunoprecipitations (IP). Non-transformed wildtype cells (WT) as control. “hit”: false discovery rate (FDR) smaller 5% and a fold-change (FC) of at least 100%; “candidate”: FDR below 20% and a FC of at least 50%. Early (about 30% fragmentation) in (B) and very late (100% fragmentation + visible new MACs) time points in (B) and (D) were collected. For DevPF1-IP (B), multiple comparisons are shown. Above each subgraph the comparison between samples is indicated, with the common condition just below. Genes of interest are labeled with one name in cases where the detected peptides cannot be unambiguously distinguished. (C) and (E): Western blots of input and IP fraction from IPs performed on DevPF1-HA (HA-affinity IP) (E) and DevPF2-GFP (GFP-affinity IP) (F) with wildtype (WT) as control. (*): These time points are not represented in the cell stage compositions in Fig S5.

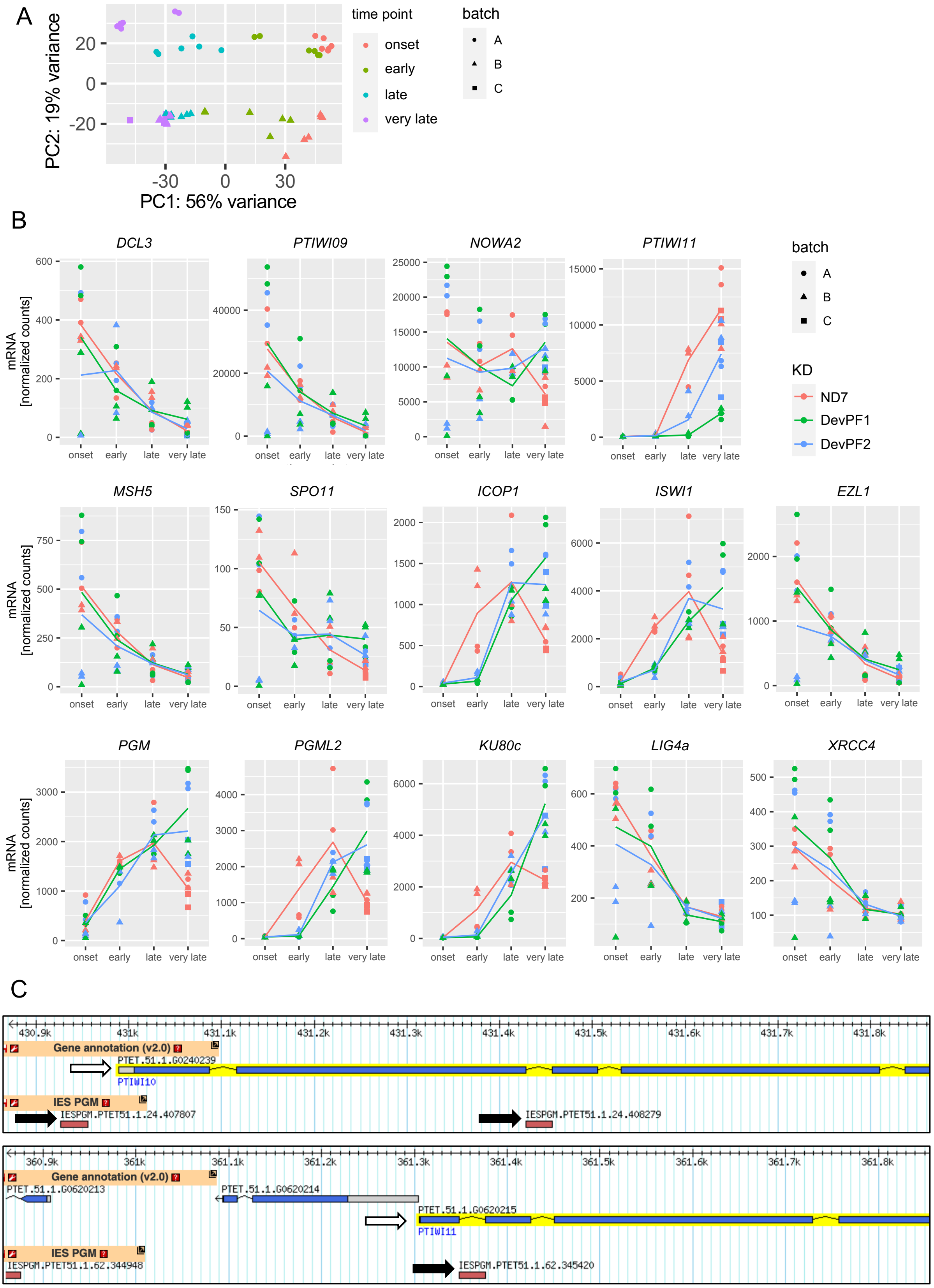
A
PC2: 19% variance
PC1: 56% variance
time point
onset
early
late
very late
batch
A
B
C
B
DCL3
PTIWI09
NOWA2
PTIWI11
MSH5
SPO11
ICOP1
ISWI1
EZL1
PGM
PGML2
KU80c
LIG4a
XRCC4
mRNA [normalized counts]
onset early late very late
batch
A
B
C
KD
ND7
DevPF1
DevPF2
C
Gene annotation (v2.0)
PTET.51.1.G0240239
PTIWI10
IES PGM
IESPGM.PTET51.1.24.407807
IESPGM.PTET51.1.24.408279
Gene annotation (v2.0)
PTET.51.1.G0620213
PTET.51.1.G0620214
PTET.51.1.G0620215
PTIWI11
IES PGM
IESPGM.PTET51.1.62.344948
IESPGM.PTET51.1.62.345420

**Figure S7: Differential expression of genes involved in *Paramecium* genome reorganization upon *DevPF* knockdowns**

(A) Principal Component Analysis (PCA) for mRNA-seq samples. (B) Gene expression levels of selected genes upon knockdowns (*ND7* (control), *DevPF1* and *DevPF2*) at developmental time points (onset, early, late and very late). The lines represent the mean of all replicates in a given knockdown and time point. (C) Screenshot from GBrowse tool on ParameciumDB showing IESs in *PTIWI10* (top) and *PTIWI11* (bottom) flanking/coding regions. White arrow: start of the gene. Black arrow: IES.

**Table S1: Lengths of the most highly retained IESs**

For each KD, the median and mean length [bp] of the top 10% most highly retained IESs are provided.

| Knockdown | Median [bp] | Mean [bp] |
|---|---|---|
| all IESs | 50 | 79.1 |
| *PGM*-KD | 67 | 130.0 |
| *KU80c*-KD | 45 | 86.8 |
| *DevPF1*-KD (2) | 66 | 143.9 |
| *DevPF2*-KD (4) | 84.5 | 134.8 |
| *SPT5m*-KD | 85 | 179.4 |
| *DCL2/3/5*-KD | 85 | 153.2 |
| *EZL1*-KD | 77 | 145.7 |
| *ISWI1*-KD | 77 | 148.6 |
| *DevPF1*-KD (1) | 62 | 124.9 |
| *DevPF1*-KD (3) | 85 | 184.1 |
| *DevPF1*-KD (4) | 75 | 160.4 |
| *DevPF2*-KD (1) | 86 | 166.9 |
| *DevPF2*-KD (2) | 76 | 141.1 |
| *DevPF2*-KD (3) | 81 | 130.0 |
| *DevPF2*-KD (5) | 79 | 133.9 |
| *DevPF2*-KD (6) | 85 | 143.6 |

**Table S2: Statistics alternative excision**

For each KD, the percentage of alternative excision events was determined for each IES.The median and mean values for all IESs are provided. Replicates are indicated in parentheses. Rep = replicates.

| Knockdown | Rep | median [%] | mean [%] |
|---|---|---|---|
| *ND7* | 3 | 0 | 2.72 |
| | 4 | 0 | 2.43 |
| *ISWI1* | - | 4.35 | 10.88 |
| *DevPF1* | 2 | 0 | 2.53 |
| | 3 | 0 | 3.82 |
| | 4 | 0 | 3.49 |
| *DevPF2* | 1 | 0 | 3.54 |
| | 3 | 0 | 3.85 |
| | 4 | 0 | 3.96 |
| | 5 | 0 | 3.52 |
| | 6 | 0 | 3.53 |

**Table S3: Protein abundance in DevPF1-IP**

Selected proteins identified by mass spectrometry in DevPF1- and DevPF2-IP. Comparison: the compared samples and, if applicable, the common condition. Protein: Ambiguous hits are labeled with one name. “hit”: false discovery rate (FDR) smaller 5% and a fold-change (FC) of at least 100%; “candidate”: FDR below 20% and a FC of at least 50%.

| Comparison | Protein | LogFC | Log(p) | Hit annotation |
|---|---|---|---|---|
| DevPF1 vs WT (early) | DevPF1 | 4.61 | 7.69 | hit |
| | Ptiwi01 | 2.29 | 7.67 | hit |
| | Histone 2B | 0.84 | 0.97 | candidate |
| | Histone 4 | 0.20 | 0.16 | no hit |
| DevPF1 vs WT (late) | DevPF1 | 3.26 | 6.00 | hit |
| | Ptiwi01 | 1.25 | 4.77 | hit |
| | Histone 2B | 2.37 | 3.49 | hit |
| | Histone 4 | 2.20 | 3.29 | hit |
| late vs early (DevPF1) | DevPF1 | -0.47 | 0.64 | no hit |
| | Ptiwi01 | -1.35 | 5.11 | hit |
| | Histone 2B | 1.60 | 2.23 | candidate |
| | Histone 4 | 1.59 | 2.26 | candidate |
| late vs early (WT) | DevPF1 | 0.89 | 1.48 | no hit |
| | Ptiwi01 | -0.30 | 0.88 | no hit |
| | Histone 2B | 0.07 | 0.05 | no hit |
| | Histone 4 | -0.42 | 0.40 | no hit |
| DevPF2 vs WT | DevPF2 | 2.67 | 6.08 | hit |
| | Ptiwi10 | 2.28 | 4.92 | hit |
| | Histone 2B | 1.70 | 1.45 | hit |
| | Histone 4 | 1.67 | 1.88 | hit |

**Table S4: Differential expression of selected genes (p-value)**

Adjusted p-values for differential expression (thresholds: |log2(Fold Change)| > 2; adjusted p-value < 0.01) of selected genes in different knockdown comparisons and time points. Magenta = upregulation; Blue = downregulation.

| | *DevPF1*-KD vs *ND7*-KD | | | | *DevPF2*-KD vs *ND7*-KD | | | | *DevPF2*-KD vs *DevPF1*-KD | | | |
|---|---|---|---|---|---|---|---|---|---|---|---|---|
| Gene | onset | early | late | very late | onset | early | late | very late | onset | early | late | very late |
| *DCL2* | 9.97E-01 | 1.00E+00 | 9.80E-01 | 6.60E-01 | 6.25E-01 | 1.00E+00 | 9.87E-01 | 9.85E-01 | 6.09E-01 | 1.00E+00 | 1.00E+00 | 1.00E+00 |
| *DCL3* | 9.96E-01 | 1.00E+00 | 9.99E-01 | 3.93E-01 | 6.85E-01 | 1.00E+00 | 9.97E-01 | 8.19E-01 | 7.26E-01 | 1.00E+00 | 1.00E+00 | 1.00E+00 |
| *DCL5* | 9.74E-01 | 9.87E-07 | 1.23E-01 | 1.97E-01 | 6.23E-01 | 7.36E-06 | 3.02E-01 | 6.29E-01 | 8.40E-01 | 1.00E+00 | 1.00E+00 | 1.00E+00 |
| *DevPF1* | 1.14E-02 | 6.41E-04 | 1.15E-01 | 5.81E-01 | 1.54E-01 | 1.00E+00 | 8.51E-01 | 9.61E-01 | 1.80E-01 | 8.07E-01 | 1.00E+00 | 1.00E+00 |
| *DevPF2* | 9.83E-01 | 6.44E-01 | 4.61E-02 | 6.63E-01 | 8.10E-01 | 2.36E-07 | 4.51E-08 | 5.54E-04 | 9.71E-01 | 2.25E-01 | 1.00E+00 | 4.75E-01 |
| *EZL1* | 9.97E-01 | 1.00E+00 | 9.20E-01 | 4.47E-01 | 6.19E-01 | 1.00E+00 | 9.53E-01 | 7.08E-01 | 6.53E-01 | 1.00E+00 | 1.00E+00 | 1.00E+00 |
| *GTSF1* | 9.93E-01 | 1.00E+00 | 9.16E-01 | 5.10E-01 | 7.25E-01 | 1.00E+00 | 9.64E-01 | 7.54E-01 | 8.26E-01 | 1.00E+00 | 1.00E+00 | 1.00E+00 |
| *ICOP1* | 9.94E-01 | 1.87E-20 | 8.06E-01 | 2.52E-03 | 6.28E-01 | 3.39E-13 | 9.97E-01 | 5.26E-03 | 5.83E-01 | 1.00E+00 | 1.00E+00 | 1.00E+00 |
| *ICOP2* | 8.98E-01 | 5.32E-25 | 9.70E-01 | 1.28E-01 | 3.22E-01 | 6.03E-14 | 7.81E-01 | 1.34E-02 | 8.69E-01 | 1.00E+00 | 1.00E+00 | 1.00E+00 |
| *ISWI1* | 8.16E-01 | 1.08E-03 | 7.46E-01 | 1.48E-02 | 9.13E-01 | 1.07E-04 | 9.90E-01 | 9.36E-03 | 3.43E-01 | 1.00E+00 | 1.00E+00 | 1.00E+00 |
| *KU80c* | 9.09E-01 | 1.44E-19 | 2.89E-01 | 5.67E-02 | 6.37E-01 | 8.69E-11 | 9.14E-01 | 4.30E-02 | 2.32E-01 | 1.00E+00 | 1.00E+00 | 1.00E+00 |
| *LIG4a* | 9.37E-01 | 1.00E+00 | 8.45E-01 | 8.04E-01 | 3.82E-01 | 1.00E+00 | 9.96E-01 | 9.92E-01 | 7.68E-01 | 1.00E+00 | 1.00E+00 | 1.00E+00 |
| *MSH4a* | 9.84E-01 | 1.00E+00 | 9.13E-01 | 7.58E-01 | 4.85E-01 | 1.00E+00 | 9.62E-01 | 9.82E-01 | 6.50E-01 | 1.00E+00 | 1.00E+00 | 1.00E+00 |
| *MSH4b* | 9.96E-01 | 1.00E+00 | 6.50E-01 | 4.96E-01 | 4.89E-01 | 1.00E+00 | 9.26E-01 | 7.73E-01 | 5.35E-01 | 1.00E+00 | 1.00E+00 | 1.00E+00 |
| *MSH5* | 9.94E-01 | 1.00E+00 | 9.75E-01 | 7.96E-01 | 7.16E-01 | 1.00E+00 | 9.94E-01 | 8.26E-01 | 7.84E-01 | 1.00E+00 | 1.00E+00 | 1.00E+00 |
| *ND7* | 1.65E-13 | 3.43E-20 | 3.25E-24 | 1.69E-46 | 2.89E-11 | 1.62E-16 | 2.85E-18 | 7.18E-51 | 6.89E-01 | 1.00E+00 | 1.00E+00 | 1.00E+00 |
| *NOWA1* | 9.87E-01 | 1.00E+00 | 8.57E-01 | 3.23E-01 | 6.00E-01 | 1.00E+00 | 9.62E-01 | 3.13E-01 | 7.21E-01 | 1.00E+00 | 1.00E+00 | 1.00E+00 |
| *NOWA2* | 9.87E-01 | 1.00E+00 | 7.92E-01 | 3.16E-01 | 5.68E-01 | 1.00E+00 | 9.60E-01 | 2.75E-01 | 7.02E-01 | 1.00E+00 | 1.00E+00 | 1.00E+00 |
| *PDSG2* | 8.53E-01 | 1.93E-13 | 6.82E-01 | 3.83E-02 | 3.18E-01 | 1.54E-12 | 9.64E-01 | 1.07E-01 | 9.93E-01 | 1.00E+00 | 1.00E+00 | 1.00E+00 |
| *PGM* | 9.31E-01 | 1.00E+00 | 9.71E-01 | 7.76E-02 | 8.30E-01 | 1.00E+00 | 9.61E-01 | 8.70E-02 | 7.76E-01 | 1.00E+00 | 1.00E+00 | 1.00E+00 |
| *PGML1* | 8.64E-01 | 1.65E-12 | 8.66E-01 | 9.71E-03 | 4.68E-01 | 2.62E-08 | 9.16E-01 | 3.97E-02 | 8.99E-01 | 1.00E+00 | 1.00E+00 | 1.00E+00 |
| *PGML2* | 9.96E-01 | 1.46E-19 | 3.47E-01 | 1.30E-02 | 8.92E-01 | 2.71E-14 | 8.99E-01 | 5.60E-03 | 9.36E-01 | 1.00E+00 | 1.00E+00 | 1.00E+00 |
| *PGML3a* | 9.49E-01 | 1.47E-12 | 9.96E-02 | 6.76E-02 | 6.66E-01 | 4.12E-08 | 9.49E-01 | 1.78E-02 | 9.95E-01 | 1.00E+00 | 1.00E+00 | 1.00E+00 |
| *PGML3b* | 8.98E-01 | 1.11E-06 | 4.02E-01 | 3.70E-01 | 9.87E-01 | 7.64E-03 | 9.11E-01 | 1.09E-01 | 5.15E-01 | 1.00E+00 | 1.00E+00 | 1.00E+00 |
| *PGML3c* | 9.87E-01 | 4.72E-01 | 4.14E-02 | 2.65E-01 | 5.12E-01 | 5.41E-01 | 4.73E-01 | 2.46E-01 | 6.44E-01 | 1.00E+00 | 1.00E+00 | 1.00E+00 |
| *PGML4a* | 9.58E-01 | 2.15E-10 | 7.33E-01 | 2.46E-04 | 8.05E-01 | 3.26E-08 | 8.67E-01 | 3.57E-02 | 9.23E-01 | 1.00E+00 | 1.00E+00 | 1.00E+00 |
| *PGML4b* | 9.20E-01 | 2.13E-08 | 9.15E-01 | 3.77E-03 | 8.67E-01 | 5.82E-07 | 8.67E-01 | 7.23E-03 | 7.08E-01 | 1.00E+00 | 1.00E+00 | 1.00E+00 |
| *PGML5a* | 9.50E-01 | 6.89E-09 | 9.28E-01 | 5.96E-03 | 5.70E-01 | 4.48E-06 | 9.26E-01 | 2.19E-02 | 9.04E-01 | 1.00E+00 | 1.00E+00 | 1.00E+00 |
| *PGML5b* | 9.94E-01 | 2.06E-17 | 4.86E-01 | 1.25E-01 | 9.25E-01 | 2.17E-09 | 9.14E-01 | 3.09E-01 | 9.89E-01 | 1.00E+00 | 1.00E+00 | 1.00E+00 |
| *PTCAF1* | 9.99E-01 | 1.00E+00 | 9.79E-01 | 4.52E-01 | 6.71E-01 | 1.00E+00 | 9.60E-01 | 6.73E-01 | 6.92E-01 | 1.00E+00 | 1.00E+00 | 1.00E+00 |
| *PTIWI01* | 9.97E-01 | 1.00E+00 | 9.65E-01 | 5.85E-01 | 8.81E-01 | 1.00E+00 | 9.79E-01 | 7.11E-01 | 8.68E-01 | 1.00E+00 | 1.00E+00 | 1.00E+00 |
| *PTIWI09* | 9.96E-01 | 1.00E+00 | 9.48E-01 | 5.78E-01 | 8.72E-01 | 1.00E+00 | 9.82E-01 | 7.56E-01 | 8.43E-01 | 1.00E+00 | 1.00E+00 | 1.00E+00 |
| *PTIWI10* | 9.77E-01 | 1.00E+00 | 6.29E-19 | 3.04E-11 | 5.72E-01 | 1.00E+00 | 7.47E-06 | 2.73E-02 | 7.70E-01 | 1.00E+00 | 1.73E-01 | 3.46E-03 |
| *PTIWI11* | 9.87E-01 | 1.00E+00 | 1.33E-17 | 1.96E-04 | 9.58E-01 | 1.00E+00 | 3.54E-03 | 4.99E-01 | 9.41E-01 | 1.00E+00 | 7.50E-03 | 2.11E-01 |
| *SPO11* | 9.77E-01 | 1.00E+00 | 8.32E-01 | 2.05E-01 | 6.36E-01 | 1.00E+00 | 9.02E-01 | 4.19E-01 | 8.36E-01 | 1.00E+00 | 1.00E+00 | 1.00E+00 |
| *SPT5m* | 9.96E-01 | 1.00E+00 | 9.66E-01 | 4.37E-01 | 6.67E-01 | 1.00E+00 | 9.81E-01 | 6.80E-01 | 7.24E-01 | 1.00E+00 | 1.00E+00 | 1.00E+00 |
| *TFIIS4* | 9.87E-01 | 1.00E+00 | 9.54E-01 | 6.85E-01 | 5.42E-01 | 1.00E+00 | 9.37E-01 | 4.74E-01 | 6.76E-01 | 1.00E+00 | 1.00E+00 | 1.00E+00 |

**Table S5: Differential expression of selected genes (fold change)**

Log2 transformed fold change for differential expression (thresholds: |log2(Fold Change)| > 2; adjusted p-value < 0.01) of selected genes in different knockdown comparisons and time points. Magenta = upregulation; Blue = downregulation.

| | *DevPF1*-KD vs *ND7*-KD | | | | *DevPF2*-KD vs *ND7*-KD | | | | *DevPF2*-KD vs *DevPF1*-KD | | | |
|---|---|---|---|---|---|---|---|---|---|---|---|---|
| Gene | onset | early | late | very late | onset | early | late | very late | onset | early | late | very late |
| *DCL2* | 0.04 | -0.28 | 0.07 | 0.60 | -0.61 | 0.27 | 0.08 | 0.04 | -0.65 | 0.56 | 0.00 | -0.55 |
| *DCL3* | -0.07 | -0.31 | 0.01 | 1.31 | -0.67 | 0.09 | 0.02 | 0.49 | -0.60 | 0.40 | 0.01 | -0.82 |
| *DCL5* | -0.23 | -2.50 | -1.15 | 0.82 | -0.43 | -2.37 | -1.09 | 0.40 | -0.20 | 0.12 | 0.07 | -0.41 |
| *DevPF1* | -2.94 | -3.13 | -1.98 | -0.70 | -1.44 | -0.88 | -0.75 | -0.10 | 1.50 | 2.25 | 1.23 | 0.60 |
| *DevPF2* | -0.16 | -0.92 | -1.38 | -0.35 | -0.19 | -2.61 | -2.74 | -1.57 | -0.03 | -1.69 | -1.36 | -1.22 |
| *EZL1* | -0.04 | -0.01 | 0.29 | 1.02 | -0.71 | -0.21 | 0.29 | 0.63 | -0.66 | -0.20 | 0.00 | -0.39 |
| *GTSF1* | -0.17 | 0.03 | 0.31 | 0.96 | -0.53 | -0.27 | 0.22 | 0.57 | -0.36 | -0.29 | -0.09 | -0.39 |
| *ICOP1* | -0.06 | -3.78 | -0.29 | 1.43 | 0.35 | -3.05 | -0.01 | 1.18 | 0.41 | 0.74 | 0.28 | -0.25 |
| *ICOP2* | -0.48 | -3.65 | -0.05 | 0.72 | -0.61 | -2.70 | 0.44 | 0.92 | -0.13 | 0.95 | 0.49 | 0.21 |
| *ISWI1* | -0.72 | -1.69 | -0.39 | 1.27 | -0.08 | -1.91 | -0.03 | 1.18 | 0.64 | -0.22 | 0.35 | -0.09 |
| *KU80c* | -0.51 | -4.13 | -0.93 | 1.13 | 0.38 | -3.08 | -0.28 | 1.07 | 0.89 | 1.05 | 0.65 | -0.05 |
| *LIG4a* | -0.37 | 0.10 | -0.28 | -0.23 | -0.60 | -0.20 | -0.01 | -0.01 | -0.24 | -0.30 | 0.26 | 0.21 |
| *MSH4a* | -0.23 | 0.01 | 0.26 | 0.43 | -0.80 | -0.27 | 0.20 | -0.05 | -0.56 | -0.28 | -0.06 | -0.47 |
| *MSH4b* | -0.03 | 0.10 | 0.68 | 0.63 | -0.62 | -0.09 | 0.30 | 0.34 | -0.58 | -0.20 | -0.38 | -0.29 |
| *MSH5* | -0.10 | -0.24 | 0.10 | 0.41 | -0.49 | -0.45 | 0.03 | 0.38 | -0.39 | -0.21 | -0.06 | -0.03 |
| *ND7* | 1.71 | 2.09 | 2.33 | 3.03 | 1.57 | 1.94 | 2.05 | 2.78 | -0.14 | -0.15 | -0.28 | -0.25 |
| *NOWA1* | -0.17 | -0.01 | -0.40 | 1.05 | -0.62 | -0.25 | -0.19 | 1.09 | -0.44 | -0.24 | 0.21 | 0.04 |
| *NOWA2* | -0.18 | -0.20 | -0.52 | 0.99 | -0.62 | -0.34 | -0.19 | 1.07 | -0.44 | -0.14 | 0.33 | 0.08 |
| *PDSG2* | -0.97 | -4.21 | -0.61 | 1.48 | -0.96 | -4.13 | 0.14 | 1.14 | 0.01 | 0.08 | 0.74 | -0.34 |
| *PGM* | -0.42 | -0.15 | 0.06 | 1.11 | -0.18 | -0.62 | 0.13 | 1.01 | 0.24 | -0.47 | 0.06 | -0.10 |
| *PGML1* | -0.74 | -3.48 | 0.25 | 1.52 | -0.60 | -2.90 | 0.28 | 1.15 | 0.13 | 0.59 | 0.03 | -0.36 |
| *PGML2* | -0.04 | -4.19 | -0.88 | 1.41 | -0.12 | -3.63 | -0.33 | 1.35 | -0.08 | 0.56 | 0.55 | -0.07 |
| *PGML3a* | -0.35 | -3.38 | -1.23 | 1.12 | -0.36 | -2.75 | -0.17 | 1.21 | -0.01 | 0.62 | 1.06 | 0.10 |
| *PGML3b* | 1.11 | -3.96 | -1.10 | 0.83 | 0.03 | -2.45 | -0.40 | 1.26 | -1.08 | 1.51 | 0.71 | 0.43 |
| *PGML3c* | 0.13 | -1.04 | -1.30 | 0.68 | 0.53 | -1.07 | -0.87 | 0.70 | 0.40 | -0.03 | 0.43 | 0.02 |
| *PGML4a* | -0.31 | -3.09 | 0.44 | 1.92 | -0.21 | -2.80 | 0.43 | 1.13 | 0.10 | 0.29 | -0.02 | -0.78 |
| *PGML4b* | -0.39 | -2.32 | 0.14 | 1.33 | -0.12 | -2.15 | 0.35 | 1.10 | 0.27 | 0.17 | 0.21 | -0.23 |
| *PGML5a* | -0.36 | -3.08 | -0.15 | 1.68 | -0.48 | -2.60 | 0.26 | 1.31 | -0.12 | 0.48 | 0.42 | -0.37 |
| *PGML5b* | -0.08 | -4.31 | -0.77 | 1.03 | -0.09 | -3.16 | -0.30 | 0.75 | -0.01 | 1.15 | 0.47 | -0.28 |
| *PTCAF1* | -0.02 | -0.22 | 0.08 | 0.94 | -0.56 | 0.01 | 0.23 | 0.65 | -0.54 | 0.23 | 0.15 | -0.29 |
| *PTIWI01* | 0.06 | -0.15 | 0.20 | 1.09 | -0.31 | -0.32 | 0.19 | 0.86 | -0.37 | -0.17 | -0.01 | -0.23 |
| *PTIWI09* | 0.11 | -0.01 | 0.30 | 1.26 | -0.38 | -0.33 | 0.18 | 0.85 | -0.50 | -0.33 | -0.12 | -0.41 |
| *PTIWI10* | -0.23 | -0.11 | -5.01 | -3.76 | -0.54 | -0.18 | -2.91 | -1.43 | -0.30 | -0.07 | 2.10 | 2.33 |
| *PTIWI11* | -0.15 | -0.72 | -5.32 | -2.61 | -0.06 | -0.18 | -2.45 | -0.73 | 0.09 | 0.54 | 2.87 | 1.88 |
| *SPO11* | -0.31 | -0.69 | 0.49 | 1.41 | -0.59 | -0.58 | 0.54 | 1.04 | -0.29 | 0.11 | 0.04 | -0.37 |
| *SPT5m* | -0.10 | -0.27 | 0.14 | 1.10 | -0.65 | -0.07 | 0.13 | 0.73 | -0.56 | 0.20 | -0.02 | -0.37 |
| *TFIIS4* | -0.17 | -0.49 | -0.13 | 0.47 | -0.64 | -0.58 | 0.29 | 0.79 | -0.47 | -0.10 | 0.42 | 0.32 |

**Table S6: Primers used in IES Retention PCRs**

| IES | Primer sequence (5' to 3' orientation) |
|---|---|
| MT Locus F | GGTGTTTATATCTTAATTGTTGACCCTCAC |
| MT Locus R | CCATCTATACTCCATTCTTTATCTTAATTCAT |
| 51G4404 F | CTGTTGCTACACATTGTGCATATGTTACT |
| 51G4404 R | GCTGTAAGATTAACATTGAGCATGATCAAG |

**Table S7: Adapter sequences used for trimming**

| IES | Illumina adapter sequence (5' to 3' orientation) |
|---|---|
| DNA-seq (read 1) | AGATCGGAAGAGCACACGTCTGAACTCCAGTCA |
| DNA-seq (read 2) | AGATCGGAAGAGCGTCGTGTAGGGAAAGAGTGT |
| mRNA-seq (read 1) | AGATCGGAAGAGCACACGTCTGAACTCCAGTCAC |
| mRNA-seq (read 2) | AGATCGGAAGAGCGTCGTGTAGGGAAAGAGTGT |
